# CRAPS: Chromosomal-Repair-Assisted Pathway Shuffling in yeast

**DOI:** 10.1101/2023.03.15.532626

**Authors:** Christien B. Dykstra, Michael E. Pyne, Vincent J.J. Martin

## Abstract

A fundamental challenge of metabolic engineering involves assembling and screening vast combinations of orthologous enzymes across a multi-step biochemical pathway. Current pathway assembly workflows involve combining genetic parts *ex vivo* and assembling one pathway configuration per tube or well. Here we present CRAPS, <u>C</u>hromosomal-<u>R</u>epair-<u>A</u>ssisted <u>P</u>athway <u>S</u>huffling, an *in vivo* pathway engineering technique that enables the self-assembly of one pathway configuration per cell. CRAPS leverages the yeast chromosomal repair pathway and utilizes a pool of inactive, chromosomally integrated orthologous gene variants corresponding to a target multi-step pathway. Supplying gRNAs to the CRAPS host activates the expression of one gene variant per pathway step, resulting in a unique pathway configuration in each cell. We deployed CRAPS to build more than 1,000 combinations of a four-step carotenoid biosynthesis network. Sampling the CRAPS pathway space yielded strains with distinct color phenotypes and carotenoid product profiles. We anticipate that CRAPS will expedite strain engineering campaigns by enabling the generation and sampling of vast biochemical spaces.

## INTRODUCTION

Metabolic engineering and synthetic biology aim to reconstruct heterologous biochemical pathways in tractable laboratory microorganisms. Major recent advances have been made in the microbial production of value-added compounds, notably artemisinin [1], benzylisoquinoline alkaloids [2], resveratrol [3], and coumaric acid [4]. The proliferation of genome and transcriptome sequencing projects has generated vast databases of genetic sequence information, such as GenBank and the thousand plant (1KP) databases [5, 6], which can be mined to compile extensive libraries of biosynthetic gene variants. In this context, optimizing a single enzymatic reaction involves synthesizing and screening dozens or hundreds of candidate orthologs, often spanning multiple domains of life. By extension, optimizing a multi-step enzyme pathway becomes a massive combinatorial challenge, as thousands of enzyme combinations must be screened. For instance, optimizing a four-step biochemical pathway consisting of 10 enzyme orthologs for each reaction yields 10^4^ or 10,000 possible pathway combinations.

Significant advancements in microbial genome editing, gene synthesis, and automated strain engineering workflows have made combinatorial strain engineering cheaper, more reliable, and more efficient. Coupling CRISPR-Cas9 genome editing with the highly efficient recombination machinery of the brewer’s yeast *Saccharomyces cerevisiae* has facilitated the development of yeast *in vivo* DNA assembly methods, such as DNA assembler [7], casEMBLR [8], and similar methods [9–11]. In organisms with less powerful homologous recombination machinery, DNA assembly methods based on DNA modifying enzymes, particularly Type IIS restriction endonucleases, have enabled the development of robust DNA assembly methods, such as BioBrick assembly, Gibson assembly, and GoldenGate cloning [12]. Still, other techniques, such as the Versatile Genetic Assembly System (VEGAS) [13], the yeast MoClo [14], and MyLo [15] toolkits, combine elements of both GoldenGate assembly and yeast homologous recombination. Furthermore, global DNA bio-foundries aim to standardize, miniaturize, and automate DNA assembly workflows [16, 17], thus liberating researchers from manual pipetting and laborious DNA manipulations. While collectively, these techniques enable sophisticated, standardized, scarless assembly and cloning of large DNA constructs, they generally demand combining genetic parts *ex vivo* and assembling one pathway configuration per tube or well.

Current methods of CRISPR/Cas9 genome editing in *S. cerevisiae* involve supplying a Cas9 and gRNA delivery plasmid along with a linear PCR-generated donor cassette for the repair of Cas9-generated double-stranded DNA breaks (DSBs). In most eukaryotic organisms, lethal DSBs can also be repaired by chromosomal homology regions, a pathway referred to as gene conversion or chromosomal repair [18]. In diploid organisms, DSBs are often repaired using the homologous chromosome as a repair template. While gene conversion is a major pathway of DSB repair in natural yeast environments, this pathway is overlooked and unutilized within the context of CRISPR-Cas9 pathway engineering, as most experiments employ haploid yeast strains or target Cas9 to both copies of homologous chromosomes in diploid strains. Previously, we exploited the yeast chromosomal repair pathway to rapidly delete seven chromosomal copies of a norcoclaurine synthase (NCS) gene using a single chromosomal repair template possessing flanking homology to the NCS gene cassette [19]. Owing to the efficiency of this approach, we set out to explore chromosomal repair as a technique for *in vivo* pathway assembly in which a single engineered strain self-assembles vast combinations of a multi-step biochemical pathway.

Here we present Chromosomal-Repair-Assisted Pathway Shuffling (CRAPS), an *in vivo* technique for assembling and shuffling multi-step biochemical pathways in a single host. We first establish the yeast chromosomal repair pathway as a viable and overlooked pathway for genome editing in *S. cerevisiae*. We probe the limits of this pathway using synthetic DNA landing pads and compare chromosomal repair to the traditional genome editing approach using exogenously supplied DNA repair templates [20]. Using these insights, we designed the CRAPS system, which utilizes a pool of inactive, chromosomally-integrated orthologous gene variants corresponding to a target multi-step pathway. Supplying gRNAs to the CRAPS host activates the expression of one gene variant per pathway step, resulting in a unique pathway configuration in each cell. We leveraged CRAPS to assemble more than 1,000 combinations of a four-step carotenoid biosynthesis network, which yielded pathway variants with distinct color phenotypes and carotenoid product profiles.

## RESULTS & DISCUSSION

### Rationale and probing of CRISPR-mediated chromosomal repair in yeast

During the construction of yeast strains harboring multiple synthetic DNA landing pads in a prior study [20], we observed highly efficient repair of Cas9-mediated DSBs in the absence of an exogenously supplied DNA repair donor. In these experiments, chromosomal landing pads possessing artificially designed homology regions to the DSB site but lacking a Cas9 target sequence were found to serve as repair templates. This activity, which we refer to as chromosomal repair (CR), is a well-known survival mechanism for repairing lethal DSBs in eukaryotic cells [18]. Owing to the seemingly high efficiency of CR in these preliminary studies, we hypothesized that CR could be leveraged as a tool for assembling and shuffling biochemical pathways in yeast.

To assess the efficiency of CR, we compared CR to the standard CRISPR-Cas9 genome editing approach, referred to herein as exogenous repair (ER), in which DSBs are repaired by exogenously supplied DNA repair templates. First, a synthetic DNA landing pad (LPA-T9-LPZ) consisting of a unique Cas9 target site (T9) flanked by 280 bp LPA and LPZ recombination arms was integrated into the yeast genome (strain LP18) at the FgF20 locus, a ‘safe-harbor’ site allowing for reliable integration of genetic elements in yeast [21–23]. ER was measured by introducing one μg of a PCR-generated repair template (LPA-REN-LPZ) along with gRNA9. In contrast, CR was measured by first integrating the same repair template (LPA-REN-LPZ) into a different chromosome in the host genome (strain LP59). The introduction of gRNA9 in a subsequent transformation guided Cas9 to the LPA-T9-LPZ landing pad, thus triggering CR (**Fig. 1a**). We measured repair viability, defined as the proportion of DSBs that were repaired, by comparing CR and ER colony counts to a control transformation using a non-targeting gRNA. In this manner, repair viabilities of 68% and 29% were achieved using one μg of ER template and one copy of the CR template, respectively (**Fig. 1b**), suggesting that ER is more effective at repairing DSBs than CR under the conditions employed. We also measured editing efficiency for both techniques, which is defined as the proportion of colonies that possess the intended editing mutation. Both editing methods were highly efficient, as editing efficiencies of 90% (CR) and 100% (ER) were obtained based on genotyping 10 random transformants.

**Fig. 1.**
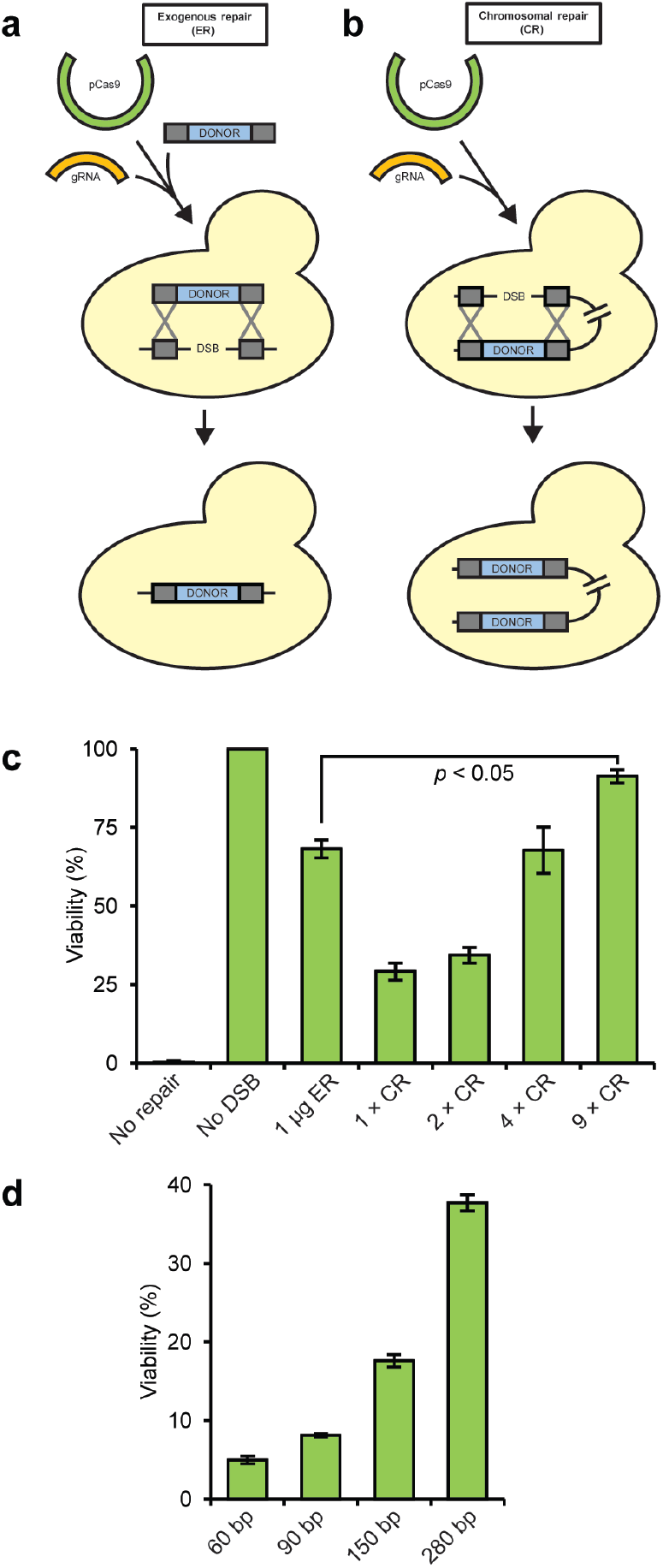
Comparison of chromosomal repair (CR) and exogenous repair (ER) pathways in yeast. **a,** Overview of ER and CR pathways. Cas9-mediated DSBs can be repaired by an exogenous or chromosomal donor. ER results in the integration of the exogenous donor, whereas CR duplicates the existing chromosomal donor. **b,** Repair efficiency of ER and CR pathways. A Cas9 DSB was introduced and repaired using one μg of an ER donor or an increasing number of CR donors. The repair template (LPA-REN-LPZ) was designed to mutate the Cas9 target site of LPA-T9-LPZ to a restriction endonuclease recognition sequence (REN) to facilitate screening. Controls are shown depicting viability in the absence of a repair template (No repair) and the absence of a gRNA (No DSB). Viability (the proportion of DSBs that were repaired) was calculated by comparing CR and ER colony counts to a control transformation using a nontargeting gRNA. Error bars represent the s.d. of three biological replicates. **c,** CR efficiency of varying-sized CR homology regions. CR templates with homology regions ranging in length from 60 bp to 280 bp were integrated into the same chromosomal locus and used to repair a Cas9 DSB. CR viability was measured as outlined in Fig. 1b. Error bars represent the s.d. of three biological replicates.

Since CR is mediated by scanning the genome for homologous repair sequences [24], we hypothesized that the efficiency of CR could be enhanced by introducing additional copies of the chromosomal repair template, thus increasing the probability of DSB repair. To test this hypothesis, we constructed strains containing two, four, or nine chromosomal repair templates (LPA-REN-LPZ) dispersed throughout the genome (strains LP7, LP8, and LP22) and compared DSB repair to the single copy CR strain (LP59) and the standard ER approach (LP18) [20]. As anticipated, CR-mediated viability improved with the number of homologous repair templates present in the yeast genome (**Fig. 1b**). By again comparing colony counts to a control transformation lacking a functional gRNA, CR in the presence of four repair templates (63% viability) was comparable to ER (72% viability). Viability using nine CR templates greatly surpassed that of ER, resulting in nearly complete repair of all chromosomal DSBs (91% viability). Genotyping of ten colonies from CR strains harboring multiple copies of LPA-REN-LPZ yielded editing efficiencies of 100% in all cases.

We also probed CR by varying the length of homology between the repair template and the DSB site (**Fig. 1c**). Repair templates were designed to possess 60 bp, 90 bp, 150 bp, or 280 bp of sequence homology (strains LP63 to LP66) and CR was assessed by repairing a single DSB with one chromosomal copy of each repair template. DSB repair was found to increase using longer homology regions. We observed more than a seven-fold increase in viability by increasing the regions of homology from 60 bp to 280 bp. Collectively, these data establish CR as a robust and efficient alternative to ER for CRISPR-Cas9-mediated genome editing in *S. cerevisiae*.

### Design of the CRAPS system

Owing to its high efficiency, we pursued CR as a tool for pathway engineering in *S. cerevisiae*. We envisioned an *in vivo* pathway assembly strategy in which CR is leveraged to build hundreds or thousands of permutations of a target metabolic network. Although our preliminary experiments utilized landing pads as CR templates, the CRAPS system utilizes genes as repair templates such that CR is coupled to the activation of gene expression. Our system consists of four synthetic Cas9 target sequences (T6, T8, T9, and T10) [20] each flanked by a different promoter and terminator pair (P_*FBA1*_-T8-T_*PRM9*_; P_*PGK1*_-T6-T_*HSP26*_; P_*PDC1*_-T9-T_*GRE3*_; P_*TDH3*_-T10-T_*TPS1*_) to facilitate high-level expression of shuffled gene variants (**Fig. 2a**). We assembled and integrated these gene-less expression cassettes into sites FgF20 (T8), FgF24 (T6), FgF18 (T9), and FgF19 (T10) of yeast (strain LP71) [20, 21].

**Fig. 2.**
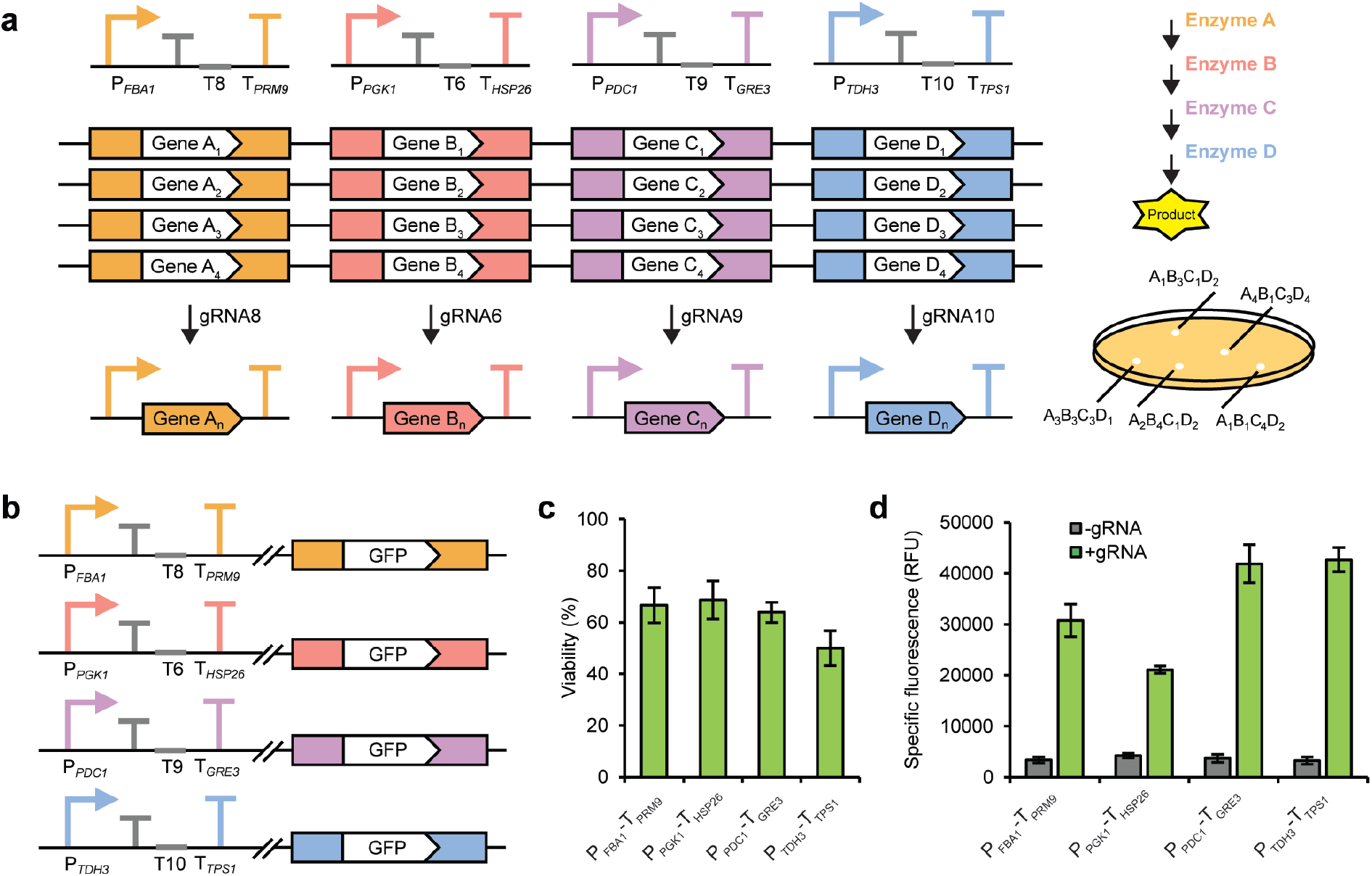
Probing the CRAPS system using a GFP repair donor and reporter enzyme. **a,** The CRAPS system consists of four gene-less expression cassettes, each possessing a unique synthetic Cas9 target site (T8, T6, T9, AND T10). A small terminator is also included downstream of each promoter to prevent transcription of Cas9 target sites. The CRAPS strain is loaded with inactive gene orthologs corresponding to a multi-step biochemical pathway. Genes possess partial homology to the flanking promoter and terminator elements (116 bp to 280 bp), rendering them inactive prior to chromosomal repair. Introduction of multiplexed CRISPR gRNAs targeting the CRAPS cassettes (gRNAs 8, 6, 9, and 10) introduces a DSB at each expression locus, activating the CR pathway and placing one gene ortholog under transcriptional control of its designated promoter. Introducing all four CRAPS gRNAs enables the self-assembly of a diverse pathway library in which each colony possesses a unique combination of orthologous pathway variants. Other combinations of two or three gRNAs also lend more pathway diversity. **b,** Construction of GFP strains for assessing CR efficiency of CRAPS cassettes. Four strains were constructed by integrating a GFP repair donor at an off-site chromosome. The *GFP* gene was designed to possess 280 bp homology to each promoter and terminator element. **c,** Assessing CR efficiency of *GFP* at each CRAPS cassette. CR viability was measured as outlined in Fig. 1b. Error bars represent the s.d. of three biological replicates. **d,** Specific GFP fluorescence (normalized to culture OD_600_) of GFP strains before and after CR induction via the introduction of gRNA. Error bars represent s.d. of three biological replicates.

To prime the CRAPS strain for pathway assembly and shuffling, pathway gene variants lacking full-length promoter and terminator elements are first dispersed throughout the genome. Variants are designed to contain only partial homology to the target promoter-terminator pair and, therefore, remain inactive prior to CR. Supplying an appropriate gRNA (gRNA6, gRNA8, gRNA9, or gRNA10) [20] introduces a DSB between the corresponding promoter and terminator, which is then repaired via host CR activity. In this manner, one random pathway variant per DSB site serves as a repair template, thus placing the selected gene under transcriptional control of the corresponding promoter-terminator pair and activating its expression. Because four promoter-terminator cassettes are present in the CRAPS strain, gene variants from up to four pathway steps can be simultaneously shuffled and expressed. The resulting pathway library comprises all theoretical pathway variant permutations and can be screened or sampled to optimize metabolic fluxes.

Before deploying CRAPS for proof-of-concept pathway shuffling, we first probed our prototype CRAPS strain for functionality. Because CRAPS relies upon partial promoter and terminator sequences to promote homology-directed repair of Cas9 DSBs, it is imperative that chromosomal gene variants remain silent prior to activation by CR. To assess this assumption, we constructed varying-sized promoter and terminator truncations of a chromosomal P_*TDH3*_-GFP-T_*CYC1*_ reporter cassette (FgF7) and quantified GFP fluorescence generated by the resulting strains (**Supplementary Fig. 1**). Mean fluorescence derived from the full-length GFP reporter cassette (653 bp P_*TDH3*_ and 250 bp T_*CYC1*_ elements) was roughly 17,000 RFU. In contrast, truncated P_*TDH3*_ promoters ranging in size from 67 bp to 250 bp generated ≤ 325 RFU. By comparison, a control strain lacking GFP produced a background mean fluorescence of 203 RFU. Based on this data, promoters of the CRAPS expression cassettes were truncated to between 116 bp to 143 bp, which coincide with predicted locations of TATA elements such that these signals were omitted from the truncated CRAPS promoters. CRAPS terminators were 280 bp in all cases.

Next, we tested the CR efficiency of our CRAPS strain at all four promoter-terminator loci by utilizing *GFP* as a repair template and reporter gene. Four strains were constructed in which *GFP* was flanked by each of the truncated promoter-terminator homology sequences and integrated at an off-target locus (strains LP75 to LP78; **Fig. 2b**). The corresponding gRNA was then introduced to each strain, triggering CR and activation of *GFP* expression. PCR screening of 12 resulting colonies from each of the four host strains revealed CR viabilities ranging from 50% to 73% (**Fig. 2c**), indicating high-level CR at each promoter-terminator locus. Mean GFP fluorescence from the four promoter-terminator pairs varied from approximately 21,000 to 43,000 RFU (**Fig. 2d**). These data demonstrate that CR can be exploited to activate high-level expression of silenced chromosomal genes by supplying Cas9 and gRNAs.

It is noteworthy that each expression cassette of the CRAPS system possesses a small transcriptional terminator [25, 26] positioned immediately downstream of the promoter element to prevent transcription of synthetic Cas9 target sites (**Fig. 2b**). Omission of these transcriptional terminators in a prior CRAPS design resulted in abnormal colony morphology and slow growth (**Supplementary Fig. 2**). Sequencing of CRAPS expression cassettes revealed varying-sized deletions in three out of the four terminator sequences downstream of the Cas9 target sites, suggesting that toxicity was associated with CRAPS expression cassettes. Redesigning the CRAPS system to prevent read-through of Cas9 target sites circumvented these deleterious issues.

#### Carotenoid pathway assembly and shuffling using the CRAPS system

To demonstrate proof-of-concept pathway assembly and shuffling using CRAPS, we turned to the carotenoid biosynthesis pathway. This pathway was selected because carotenoids are intensely colored end products that accumulate to appreciable quantities in microorganisms, enabling simple detection and qualitative comparison of carotenogenic strains. Further, the carotenoid biosynthesis network is hierarchical and highly branched, owing to the inherent promiscuity exhibited by many carotenoid biosynthetic enzymes. Several carotenoid enzyme variants have been generated through directed evolution [27–30], thus expanding the range of colored products that can be produced.

We designed a four-step carotenoid biosynthesis network for pathway assembly and shuffling using CRAPS **(Fig. 3a)**; more than 15 heterologous carotenogenic gene variants that enable a broad spectrum of colored end products were compiled, synthesized, and integrated into seven safe-harbor loci in the CRAPS strain for shuffling (**Fig. 3b**) [11, 21–23, 31]. Our carotenoid biosynthesis network incorporates overexpression of endogenous yeast genes (*tHMG1* and *BTS1*) that encode potentially rate-limiting enzymes within the yeast isoprenoid and carotenoid precursor pathways [32, 33], as well as duplicate copies of heterologous *crtI* gene variants encoding phytoene desaturase, a potentially rate-limiting enzyme in carotenoid biosynthesis [33]. The yeast isoprenoid pathway supplies geranylgeranyl pyrophosphate (GGPP), the last native carotenoid precursor, which is converted to phytoene, a colorless carotenoid. Accordingly, our background carotenoid CRAPS strain harbors chromosomal copies of GGPP synthase from *Xanthophyllomyces dendrorhus (crtE)* and phytoene synthase (*crtB*) from *Pantoea ananatis* for constitutive production of phytoene. By deploying CRAPS combinatorics using our pool of chromosomal pathway variants, a diversity of colored carotenoid products can be synthesized from phytoene.

**Fig. 3.**
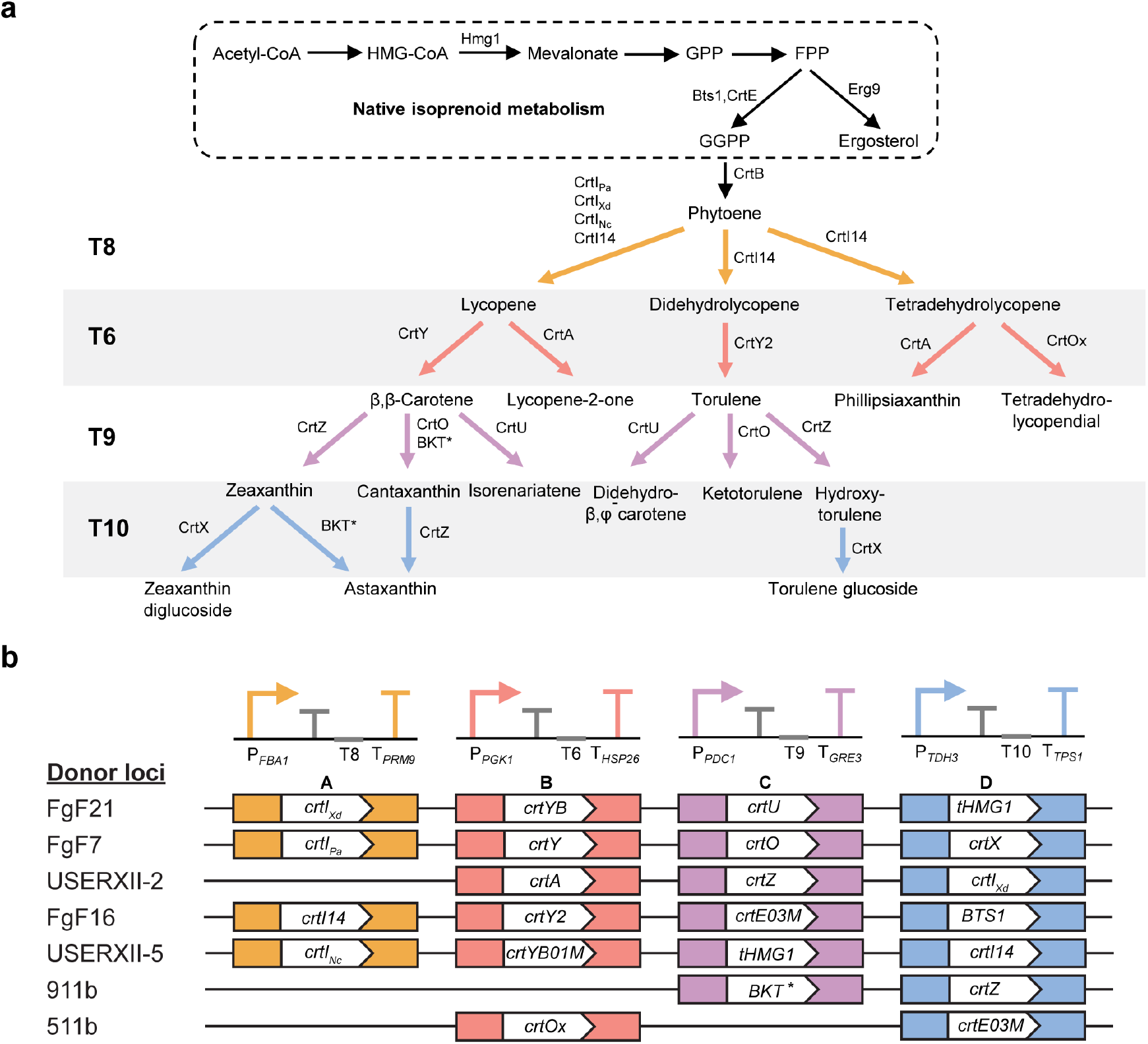
Design of a 1,000-member carotenoid biosynthesis network using CRAPS. **a,** The biosynthesis of carotenoids begins with the conversion of acetyl-CoA to farnesyl pyrophosphate (FPP) via geranyl pyrophosphate (GPP) in the yeast isoprenoid biosynthesis pathway. The intermediate FPP is a precursor to ergosterol biosynthesis or is converted to geranylgeranyl pyrophosphate (GGPP), a precursor to the heterologous carotenoid pathway. Phytoene, the committed carotenoid precursor, is synthesized from GGPP by phytoene synthase (CrtB) from *Pantoea ananatis*. Phytoene is converted to lycopene or its derivatives by any of the selected CrtI variants. Subsequent branched downstream enzymatic steps produce a variety of colored carotenoid compounds. Arrow colors indicate the target locations of the carotenoid pathway variants. **b,** Identities of all 23 carotenogenic genes loaded into the CRAPS strain for assembling a 1,000-member carotenoid biosynthesis network. Carotenogenic genes are color coded according to the corresponding CRAPS expression cassette and the carotenoid biosynthesis network outlined in **a**. Genes were integrated into chromosomes of *S. cerevisiae* in seven successive transformations.

### Multiplexed gRNA delivery and pathway assembly

To target up to four CRAPS expression sites in concert, we adapted the GTR-CRISPR method for multiplexed delivery of gRNAs [34]. Using this approach, transcription is initiated by P_SNR52_, gRNA sequences downstream of gRNA8 are preceded by a tRNA_gly_ element to facilitate gRNA processing, and gRNA arrays are terminated by T_SNR52_. In addition to targeting all four CRAPS loci in a single transformation using all four gRNAs (gRNA8, gRNA6, gRNA9, gRNA10), further combinatorial possibilities can be achieved by introducing combinations of one, two, or three gRNAs. We selected gRNA combinations based on substrate compatibility with subsequent enzymatic steps; for instance, T9 enzymes require precursors from T6 enzymes. We envisioned six relevant gRNA combinations, which we designated series 0 (gRNA8), series 1 (gRNA8, 6, 9, and 10), series 2 (gRNA8, 6, and 9), series 3 (gRNA8 and 6)), series 4 (gRNA8 and 10), and series 5 (gRNA8, 6, and 10) (**Supplementary Fig. 3**).

To verify the integration efficiency of multiplexed chromosomal repair, we introduced all six gRNA combinations in separate transformations (series 0 to 5) and randomly selected colonies for genotyping of the corresponding CRAPS loci (**Fig. 4a)**. Series 0, 3, 4, and 5 demonstrated 100% integration efficiency across all target loci (*n* = 50, 37, 53, and 50 colonies, respectively). Series 1 and 2 showed minor decreases in integration efficiency: targeting the T6 and T10 loci of series 1 yielded integration efficiencies of 92% and 83%, respectively (*n* = 24 colonies each), while targeting the T8, T6, and T9 loci of series 2 generated integration efficiencies of 93%, 95%, and 98%, respectively (*n* = 36 colonies each), demonstrating efficient multiplexed gRNA targeting and CR efficiency using up to four gRNAs (**Fig. 4a**).

**Fig. 4.**
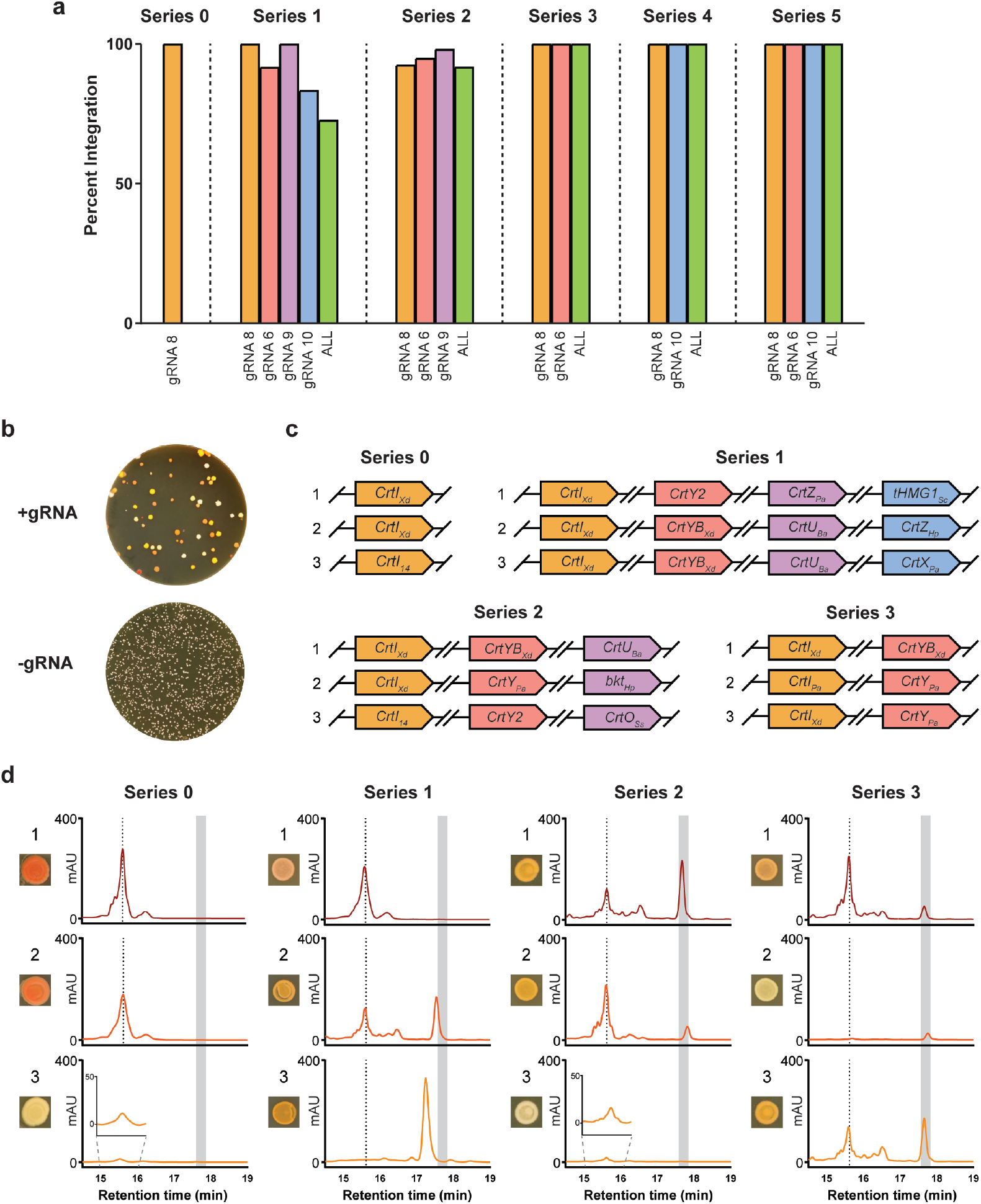
Assembling a 1,000-member CRAPS pathway library. **a,** Targeting efficiency of the four CRAPS loci performed with their respective plasmids. Integration events are determined per locus or per colony, where ‘all’ denotes that all the targeted loci were successfully converted in a single colony. Results are expressed as a percentage of the total number of transformants screened. **b,** Demonstration of a CRAPS transformation along with a control plasmid lacking targeting gRNAs. No coloration was observed in the negative control transformants. **c,** Genotypes of colonies selected for genotypic and phenotypic analysis. Genotypes were determined for colonies labeled from 1 to 3 from each of series 0 (single T8 gRNA), series 1 (all four gRNAs), series 2 (T8, T6, and T9 gRNAs), and series 3 (T8 and T6 gRNAs). **d,** High-Performance Liquid-Chromatography coupled to UV/Vis spectroscopy was performed on the same three colonies from **c**. The retention times (minutes) of peaks were recorded at 450 nm with a maximal range of 500 relative absorbance units (mAU). The dotted line represents the tentative lycopene peak, and the gray bar shows the retention time of an authentic β-carotene standard.

colonies with a diversity of carotenoid color phenotypes. Several transformants showed deep coloration in comparison to the control strain (pCD1; no gRNA), which had no discernable pigmentation (**Fig. 4b**). Comparison of the number of colony-forming units (CFUs) of CRAPS transformations using different gRNA combinations revealed decreases in viability with increasing gRNA-induced DSBs, which required scaling the CRAPS transformations accordingly to attain comparable numbers of CFUs (**Supplementary Fig. 4**).

### Sampling the CRAPS pathway space

Following multiplexed CRISPR targeting with chromosomal donors, we sampled carotenoid-producing CRAPS colonies for genotypic and metabolic analysis. For each gRNA combination series, we picked colonies with clear differences in morphology and pigmentation to highlight CRAPS’s utility in generating diverse pathway configurations and phenotypes (**Supplementary Fig. 5**). Using PCR and Sanger sequencing, we genotyped four colonies from each gRNA series (**Supplementary Table 1**), which revealed a preference for specific carotenoid gene repair donors. Specifically, *crtI_Xd_*, which was harbored at the FgF21 locus, represented 70% of integrated genes at CRAPS locus T8 across all six gRNA combination series (*n* = 24 colonies, **Supplementary Table 1, Supplementary Fig. 6**). Carotenoid gene donors integrated at the FgF21 locus (*crtIXd*, *crtYB*, *crtU*, *tHMG1*) were the most abundant repair donors at all four CRAPS loci. Conversely, we did not observe chromosomal repair of any carotenoid genes from the USER12-5 chromosomal locus (*crtI_Nc_*, *crtYB01M*, *tHMG1_Sc_*, and *crtI14*) following genotyping of 24 random colonies. Other genes not utilized for repair were *crtOx_Sa_* (511b) at the T6 locus (*n* = 16) and *crtE03M* (FgF16) at the T9 locus (*n* = 8) (**Fig. 4b, Supplementary Table 1**). We suspect that these chromosomal repair biases result from the genomic locations of the chromosomal donor templates, as none of the donors integrated at the USER12-5 site were used for repair, whereas donors in locus FgF21 facilitated efficient repair of all four CRAPS target sites (**Supplementary Fig. 6**). It is likely that the limited sample size of genotyped colonies (*n* = 8-24 colonies per locus) did not enable an accurate assessment of chromosomal site biases. It is important to note that these biases may also be partly attributed to preferentially selecting diverse colors and colony shapes. For instance, series 0 colonies expressing the fungal *crtI_Xd_* variant yielded intensely colored colonies despite comprising only a single enzymatic step from phytoene. Alternatively, screening a larger subset of colonies will provide a better comparison of chromosomal loci for gene conversion efficiency. There may be explanations for biases not attributable to experimental design that might allow for tuning of the system, such as the GC-content of donor homology regions [35]. Additionally, there exists the possibility that certain pathway configurations lead to toxic products in yeast, although this is not likely; most of the carotenogenic genes used in this study had been previously engineered into *S. cerevisiae*, and those that had not been reported did appear in viable CRAPS colonies (i.e., *crtO_Ss_*, *crtI14*, and *crtX_Pa_*, **Supplementary Table 1**) Most importantly, however, we did not observe repeated gene combinations for gRNA series targeting two or more CRAPS loci (**Supplementary Table 1**), demonstrating effective pathway assembly and shuffling using the CRAPS system.

Given the utility of CRAPS for generating genotypic diversity, we next asked if our carotenoid pathway library yielded a diversity of carotenoid metabolites. We performed HPLC analysis on carotenoid-producing strains to link product profile to pathway genotype (**Fig. 4c and d, Supplementary Table 1**). As mentioned, series 0 transformants (gRNA8) showed a bias towards *crtI_Xd_*, and two series 0 *crtI_Xd_* strains assayed by HPLC showed a chromatographic peak with a retention time of 15.6 minutes (colonies 0-1 and 0-2), whereas a colony that integrated *crtI_14_* (colony 0-3) had a reduced amount of the same peak. In addition to lack of other potential products from this enzyme, namely didehydrolycopene and tetradehydrolycopene, we presume poor catalytic efficiency in yeast of this engineered bacterial CrtI14 variant relative to the fungal CrtI_Xd_ ortholog in *S. cerevisiae*. The 15.6 minute peak is consistent with lycopene (**Fig. 4d**; dotted line), as it is the first product in our CRAPS pathway network, and CrtI_Xd_ has been shown to catalyze the conversion of phytoene to lycopene in *S. cerevisiae* [33] (**Fig. 3a**). Indeed, most of the samples that were analyzed showed the presence of this peak at 15.6 minutes (**Fig. 4d, Supplementary Fig. 7**), and spectral scanning showed absorbance maxima of 443 nm, 471 nm, and 502 nm consistent with that of lycopene in previous reports (**Supplementary Fig. 8**) [36, 37].

While cultures of series 0 (gRNA8) *crtI_Xd_* strains produced a peak consistent with lycopene, the introduction of gRNA6 in series 3 (gRNA8 and 6) facilitated the synthesis of β-carotene based on comparison to an authentic β-carotene standard. Cultures of colony one from series 3 (colony 3-1) produced appreciable amounts of the tentative lycopene peak due to the *crtI_Xd_* variant, with a small amount of β-carotene through the action of *crtYB_Xd_*. The combination of *crtI_Pa_* and *crtY_Pa_* (colony 3-2) was unfavorable for the production of lycopene and/or β-carotene, which was likely due to poor activity of CrtI_Pa_ because lycopene production was restored in colony 3-3 via the substitution of CrtI_Xd_.

Addition of gRNA9 in series 2 (gRNA8, 6, and 9) enabled the assembly of a three-step carotenoid pathway from phytoene. The T9 locus of series 2, colony one (colony 2-1) harbored a carotenoid χ-ring synthase (*crtU_Ba_*), although only tentative lycopene and β-carotene peaks were detected due to the presence of *crtI_Xd_* and *crtYB*. The T9 locus of colony 2-2 possessed a ketolase, *BKT_Hl_*\*, which converts β-carotene to canthaxanthin (**Fig. 3a, Supplementary Tables 1 and 5**). We observed a minor shift in retention time that is potentially a result of BKT_H1_* activity in concert with CrtI_Xd_ and CrtY_Pa_ (**Fig. 4b**). Colony 2-3 did not show a peak corresponding to β-carotene, again presumably due to a deficiency of CrtI14 in producing the precursor lycopene (**Fig. 4c and d**).

Cultures of CRAPS colonies transformed with all four gRNAs (series 1; gRNA8, 6, 9, and 10) produced distinct carotenoid HPLC peaks not observed in other gRNA combination series (**Fig. 4d; Supplementary Fig. 7**). Colony 1-2 (*crtI_Xd_*, *crtYB*, *crtU_Ba_*, and *crtZ_Hp_*) expressed both a χ-ring synthase (*crtU_Ba_*) and a β-carotene hydroxylase (*crtZ_Hp_*), which catalyze formation of isorenieratine/renierapurpurin or zeaxanthin, respectively (**Fig. 4c and d**). Cultures of colony 1-1 (*crtI_Xd_*, *crtY2*, *crtZ_Pa_*, and *tHMG1_Sc_*) produced a single peak consistent with lycopene, which is likely due to an inability of CrtY2 to convert lycopene to β-carotene in *S. cerevisiae* [38] (**Fig. 3a**).

In addition to 450 nm chromatograms, we analyzed chromatograms recorded at 260 nm and did not observe the production of unique peaks in comparison to controls and spectral analysis across all samples confirmed these species were likely the same (**Supplementary Fig. 8**). Finally, it is worth noting that we did not observe carotenoid peaks at 450 nm that suggest leaky expression of the silent CR donors, as the carotenoid CRAPS strain failed to synthesize carotenoids in the absence of gRNAs (**Supplementary Fig. 7 and 8**). Collectively, our genotypic and phenotypic results demonstrate the ability to rapidly generate pathway and metabolite diversity using the CRAPS system.

### Conclusion

Owing to the proliferation of genome and transcriptome sequencing databases, combinatorics has become a major challenge in synthetic biology and metabolic engineering. Our CRAPS system is capable of rapidly and efficiently generating vast combinatorial pathways, which can streamline strain optimization for various metabolic engineering goals. Using CRAPS, we have demonstrated the construction of many diverse carotenogenic strains of *S. cerevisiae* and confirmed the functional expression of these pathways based on the synthesis of a diversity of carotenoid products. CRAPS addresses many of the challenges associated with combinatorial pathway assembly, as our method allows for a single strain to generate myriad combinations in a one-step transformation, bypassing issues involved with episomal plasmid library preparation and integration of large exogenously supplied pathways. Furthermore, the use of chromosomal integration enables genotypic stability without the need for antibiotic or auxotrophic selection. Lastly, CRAPS represents a valuable genetic tool that may also be applied beyond the scope of metabolic pathways, such as deploying combinatorial gRNA libraries, assembling heterologous protein complexes, or querying transcriptional regulator function.

## METHODS

### Strains and growth media

The quadruple-auxotroph *S. cerevisiae* strain CEN.PK2-1D was used to build the CRAPS strain (LP71). Cultures were grown in YPD medium alone (10 g L^-1^ Bacto Yeast Extract, 20 g L^-1^ Bacto peptone, 20 g L^-1^ glucose), or supplemented with 200 μg mL^-1^ G418 or hygromycin B to maintain transformed plasmids. For plasmid isolation and amplification, *E. coli* DH5-α was grown in LB Miller medium (LB, Millipore) with ampicillin or carbenicillin (100 μg mL^-1^).

### Chemical transformation

Yeast transformations were carried out using an adapted version of the Gietz method [39]; Overnight yeast culture grown to an OD_600_ of ~1.0 (2 mL) was pelleted and washed with distilled water followed by 100 mM LiOAc solution. Pellets were then resuspended in 150 μL transformation solution (PEG, 33.3%; lithium acetate, 100 mM; denatured salmon sperm DNA 300 ng μL^-1^) containing 1 μg of targeting plasmid and/or 500 – 2000 ng of linear donor DNA. Samples were heat-shocked at 42°C for thirty minutes prior to 24-hour recovery in 600 μL of YPD broth. Transformants were selected using media containing geneticin (G418, 200 ng μL^-1^) and cured on YPD media.

### Yeast strain construction

A complete list of plasmids, constructs, and integration targets used in this study can be found in the supplementary information (**Supplementary Tables 2 to 5)**. For multiplexed gRNA plasmid construction, multi-part constructs were generated using Golden Gate assembly (NEB).

Purification of plasmids was done with the Thermo Scientific GeneJET Plasmid Miniprep Kit (Thermo Fisher Scientific). The pCAS vectors with split-G418 selection were adapted from Bean *et al*. [15]. The insertion of simple constructs such as single-target gRNA expression systems and resistance cassettes was completed either by ligating homologous overhangs or homology-directed repair *in vivo*. To assemble a single or multiple gRNAs into the pCAS vector, oligonucleotides containing protospacer sequences and gRNA scaffold homologies were used to amplify parts for two-, three-, or four-part golden gate assembly (**Supplementary Fig. 9**). Prior to transformation, pCAS vectors were linearized with AatII and SbfI at a region with an inactive G418 resistance marker bearing an internal 500bp homology region to allow for *in vivo* repair. Only those yeasts that successfully re-circularized the vector were resistant to G418 [15] (**Supplementary Fig. 9**). Integration of donor cassettes with flanking homology regions was performed using CRISPR/Cas9 with gRNAs specific to each safe-harbor locus [20]. A complete list of primers used for plasmid and part amplification can be found in the supporting information (**CRAPS Primers**).

### Carotenoid extraction

CRAPS-transformed strains and controls were grown in 10 mL of yeast-peptone dextrose media for 48 hours at 30 °C to similar OD_600_ values, and 5 mL of each culture was washed and pelleted. Following a method previously described by Xie *et al*. [40], hydrochloric acid (3N) was added to the samples, followed by vigorous vortexing before placing them in a hot water bath (98 °C) for 5 minutes to lyse the pellets. Samples were placed immediately on ice, and lysates were pelleted at high speed. To extract the carotenoids, acetone was added to the pelleted debris, followed by vortexing and incubation at 50 °C for 10 minutes. This process was repeated three times to remove as much coloration as possible (800 μL of acetone total), and the final extracts were clarified by spinning at max speed for 10 minutes before HPLC analysis. Each series was extracted and analyzed independently.

### HPLC analysis of metabolites

We adapted a method by Rivera and Canela for rapid carotenoid separation and qualification [41]. HPLC-PDA was performed using an Agilent 1200 system equipped with a G1315C SL diode array detection (DAD) module (Agilent, Germany). DAD spectrum scanning was from 210 nm to 550 nm, and chromatograms were recorded at 260 nm and 450 nm. Agilent ChemStation software was used for instrument control and data acquisition. Reverse-phase chromatography was carried out using an ACQUITY UPLC BEH 130A C18 column, 2.1 mm × 50 mm (Waters) maintained at 30 °C. A mobile phase gradient system consisting of solvent A, acetonitrile-methanol (7:3, v/v), and solvent B, H_2_O (100%) was used; The method begins with 80:20, A:B, followed by a gradient beginning from 2:40 to 12:00 where A is increased to 100%. After 18:00, the column is washed with 80:20, A:B, before beginning the next run. The sample injection volume was 10 μL, and each run was preceded by a 3-minute wash of 100% solvent A. A β-carotene standard, 100 μM, dissolved in an extraction matrix was used as a reference for comparing the diversity of peaks obtained.

### Statistical analyses

All numerical values are depicted as means ± s.d unless stated otherwise. Statistical differences between control and derivative strains were assessed via two-tailed Student’s t-tests assuming equal variances.

## Supporting information

Primer List

