## Supplemental Information for "CRAPS: Chromosomal-Repair-Assisted Pathway Shuffling in yeast"

### Supplementary Results and Figures

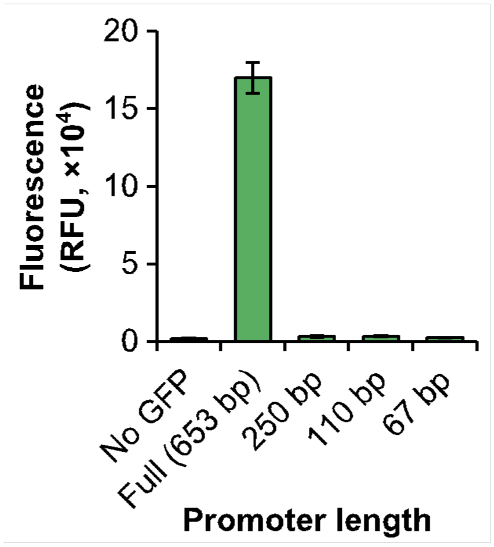


**Supplementary Figure 1. GFP fluorescence of yeast cultures harboring truncated P*TDH3* promoters.** *GFP* was expressed from various-sized P*TDH3* truncations (250 bp, 110 bp, and 67 bp) and fluorescence was measured and compared to a control strain harboring full-length P*TDH3* (680 bp) and a control strain lacking GFP.

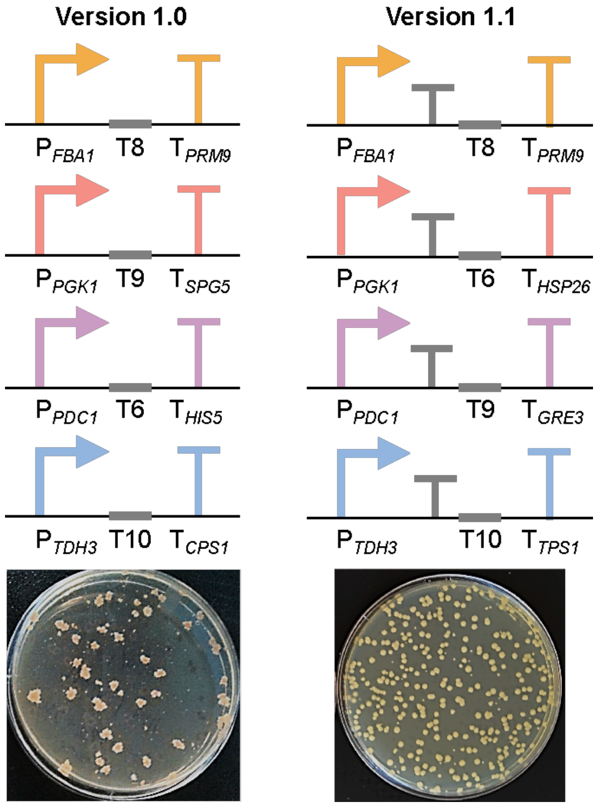


**Supplementary Figure 2. Transcription of Cas9 target sites impairs yeast growth in an early CRAPS prototype.** The omission of transcriptional terminators upstream of synthetic Cas9 target sites in an early CRAPS prototype (version 1.0) led to abnormal colony morphology and impaired growth. The subsequent CRAPS host reported herein (version 1.1) incorporates small transcriptional terminators immediately downstream of CRAPS promoters to prevent transcription of Cas9 target sites. Inclusion of such terminators circumvented growth issues associated with our first-generation CRAPS strain.

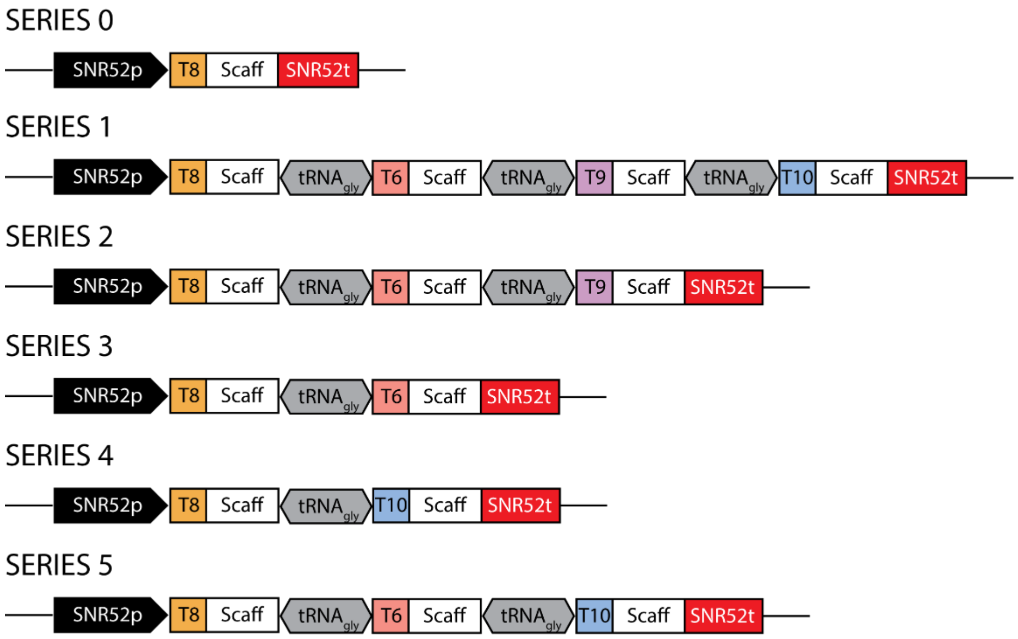


**Supplementary Figure 3. CRAPS gRNA expression series.** Multiplexed gRNA expression systems are adapted from Zhang *et al.* and delivered by pCD3 and pCD5-pCD9 [1]. To deliver a single gRNA (series 0), $P_{SNR52}$ drives transcription of the CRISPR protospacer and scaffold, which are terminated by $T_{SNR52}$. For multiplexed gRNA expression, guide-scaffold sequences are followed by $tRNA_{gly}$. Series 1 expresses all four targeting gRNAs. Series 2 targets the T8, T6, and T9 loci. Series 3 targets the T8 and T6 loci while series 4 targets T8 and T10, bypassing T6 and T9. Series 5 targets T8, T6, and T10.

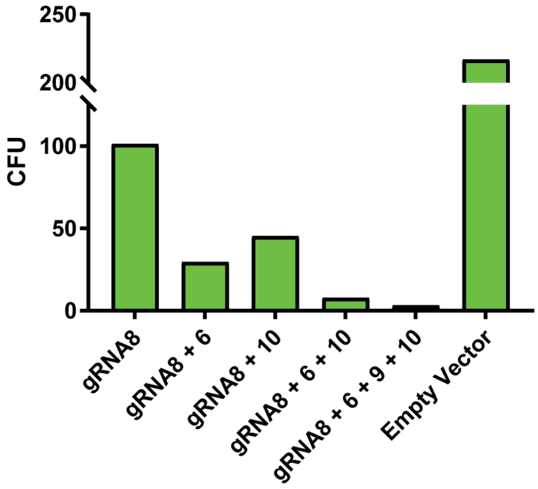


**Supplementary Figure 4. Increasing the number of double-stranded breaks introduced to the CRAPS strain decreases viable colony counts.** The viability of the CRAPS strain drops as the number of Cas9-induced double-stranded breaks increases from 1 to 4. Colony-forming units were calculated per 1 μg of empty or targeting plasmid (pCD1, pCD3, pCD5-pCD9).

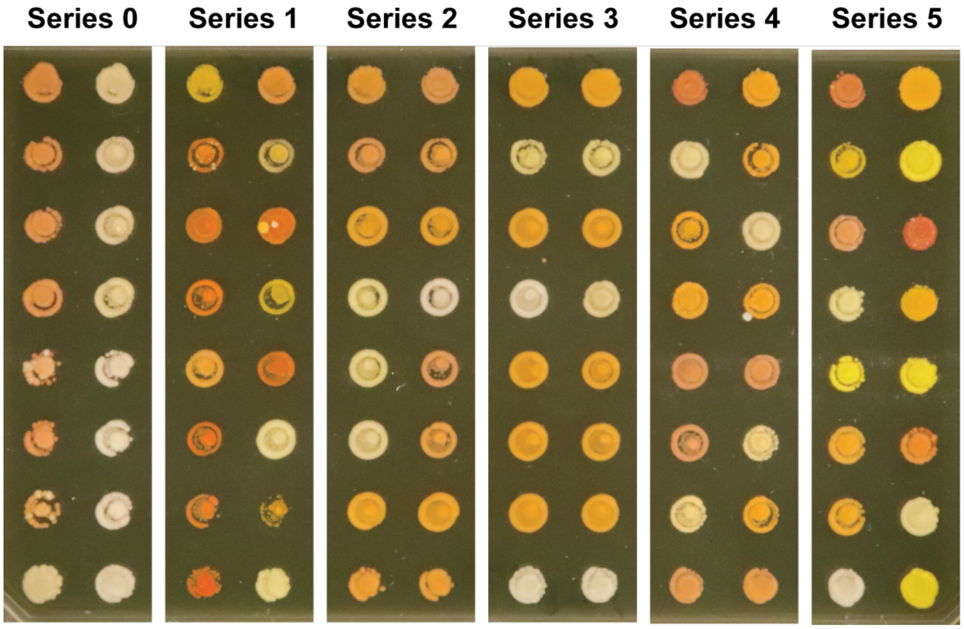


**Supplementary Figure 5. Screening plate of randomly selected colonies.** Colonies were randomly selected using a colony picker or by plating dilutions and selecting every colony on a single plate. Subsamples from each series were selected, screened for integration efficiency, and analyzed for metabolite production.

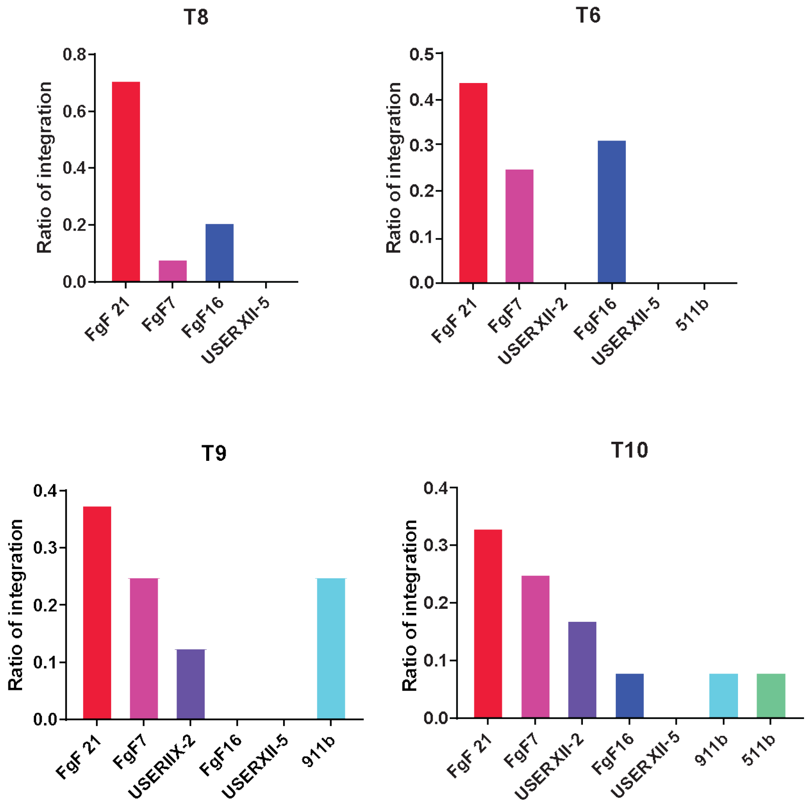


**Supplementary Figure 6. Gene conversion frequency between target and donor loci.** Each of the four CRAPS target loci (T8, T6, T9, and T10) was assessed for DSB repair frequency based on the chromosomal loci of carotenogenic gene donors.

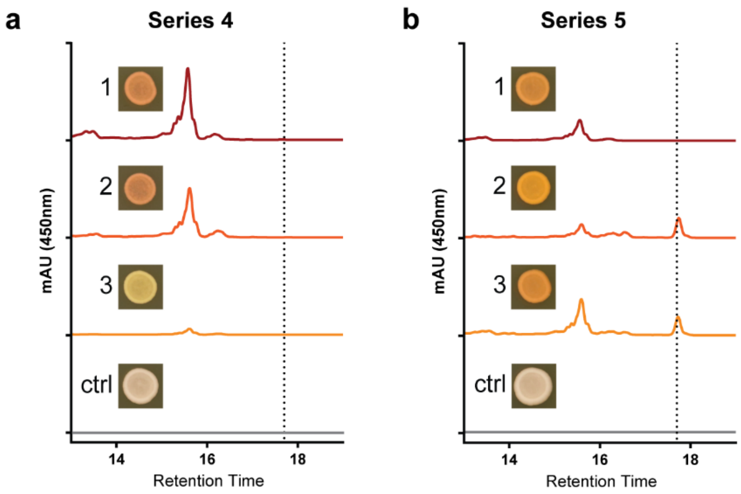


**Supplementary Figure 7. Supplementary HPLC chromatograms. a.** Chromatograms from series 4 where T8 and T10 were targeted; sample 1 incorporated *crtI*$_{Pa}$ and crtI$_{Xd}$. Sample 2 incorporated *crtI*$_{Xd}$ and *tHMG1*$_{Sc}$. Sample 3 incorporated *crtI14* and *tHMG1*$_{Sc}$. **b.** Chromatograms from series 5 where T8, T6, and T10 were targeted; Sample 1 incorporated *crtI*$_{Xd}$, *crtY2*, and *crtI*$_{Xd}$. Sample 2 incorporated *crtI*$_{Xd}$, *crtYB*$_{Xd}$, and *crtX*$_{Pa}$. Sample 3 incorporated *crtI*$_{Xd}$, *crtYB*$_{Xd}$, and *tHMG1*$_{Sc}$. Both controls were transformed with a no-gRNA control vector (pCD1). All series' chromatograms are scaled from 0 to 500 mAU at 450 nm absorbance. The dotted line corresponds to authentic β-carotene standard.

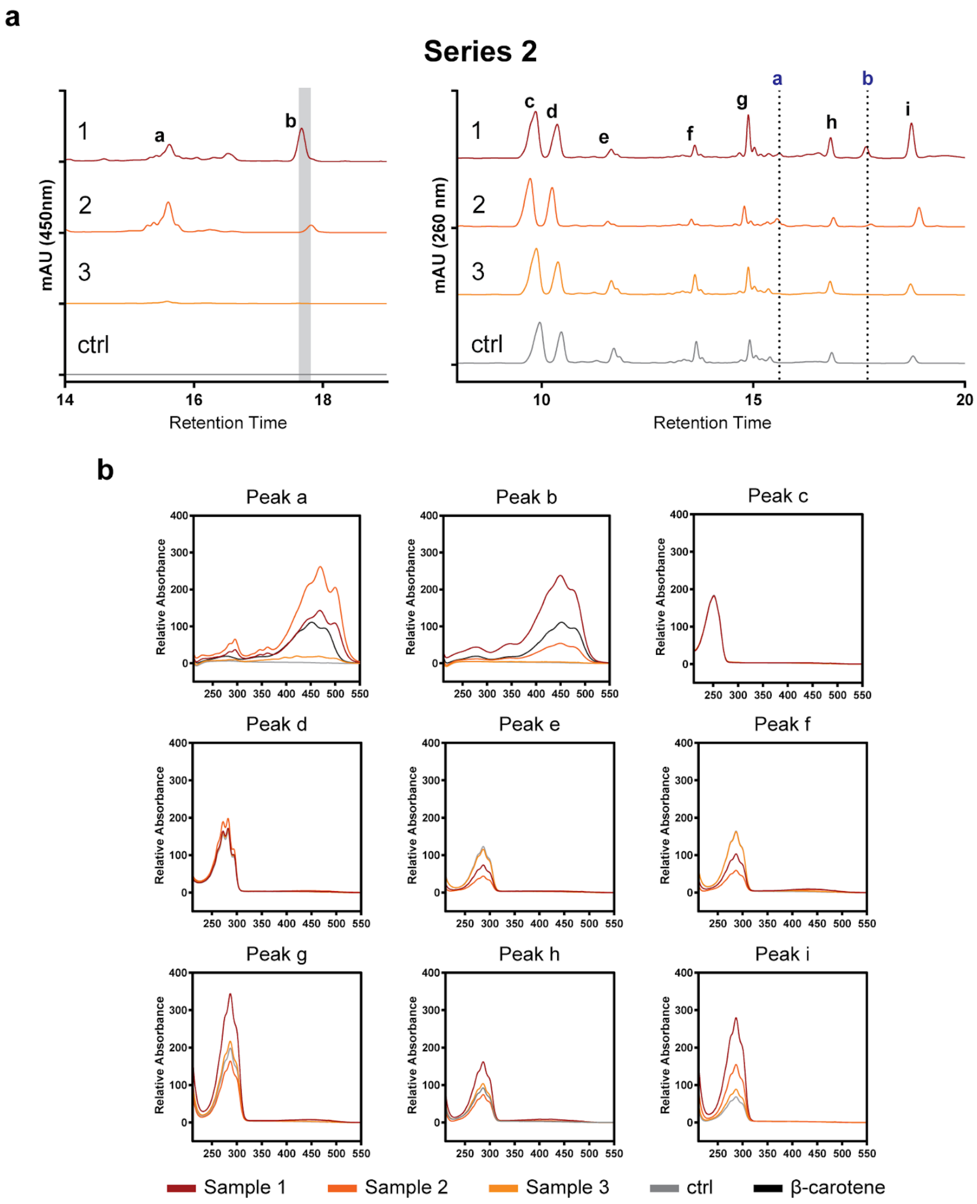


**Supplementary Figure 8. Spectral scans of peaks encountered (series 2). a.** Chromatograms from series 2 samples scanned at 450 nm (range = 500 mAU) and 260 nm (range = 200 mAU). The colony genotypes are as follows: sample 1 incorporated $CrtI_{Xd}$, $CrtYB_{Xd}$, and $CrtU_{Ba}$; sample 2 incorporated $CrtI_{Xd}$, $CrtY_{Pa}$, and $bkt_{Hp}$; sample 3 incorporated *CrtI14*, *CrtY2*, and $CrtO_{Ss}$. The grey chromatograms are from no-gRNA controls (pCD1). Dotted lines represent the retention times of peaks identified at 450 nm, and the grey bar represents authentic β-carotene standard. **b.** Absorbance spectra of identified peaks. Scans were performed at the same retention for all samples and controls and spectra were overlaid. β-carotene standard spectra were included for peaks 'a' and 'b'.

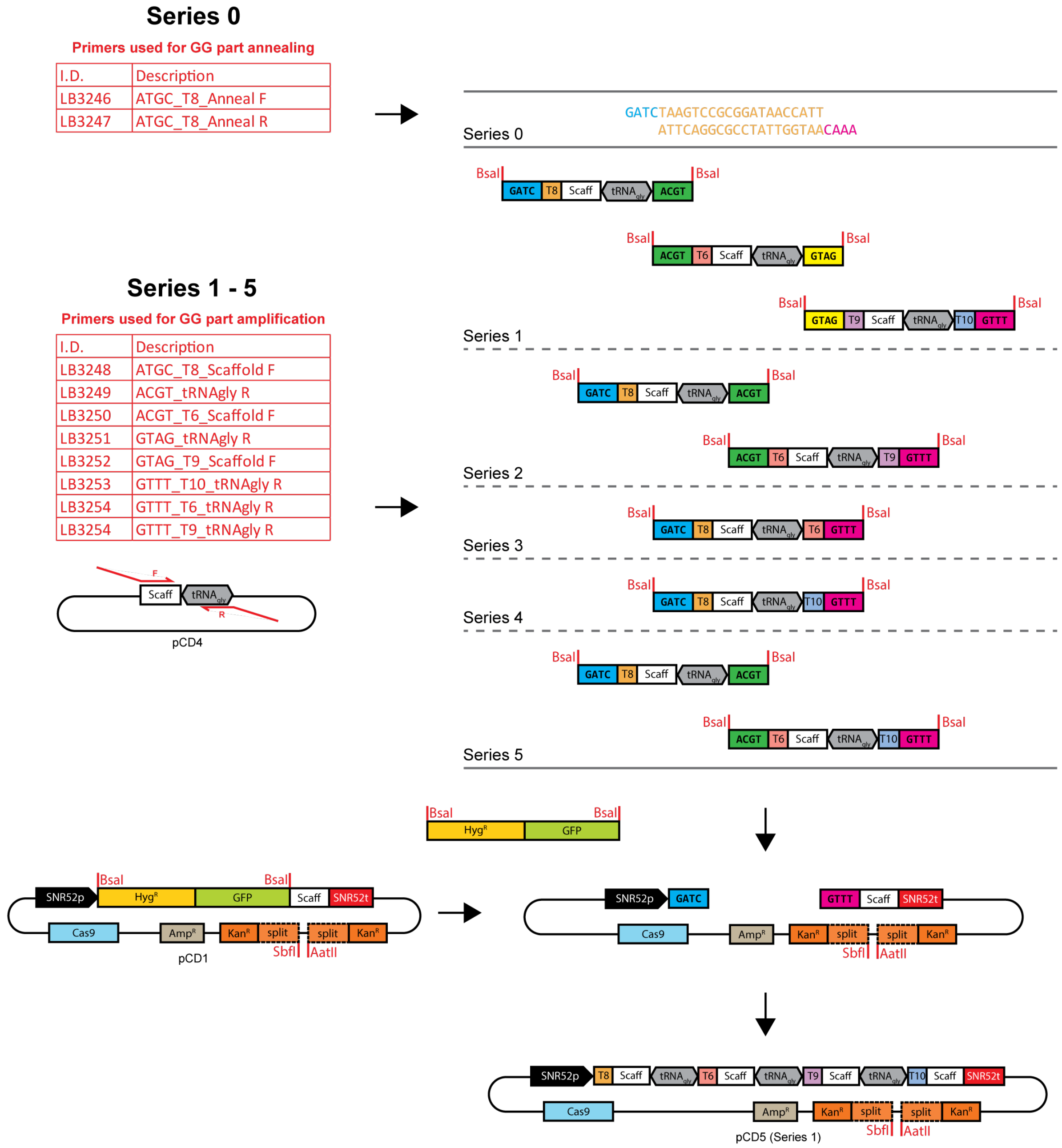


**Supplementary Figure 9. Golden Gate assembly of multiplexed gRNA targeting vectors**. Strategy adapted from Zhang *et al.* and constructed with Golden Gate Assembly [1, 2]. For the single-gRNA group (series 0), primers with Golden Gate overhangs were annealed. For multi-gRNA constructs (series 1 to 5), parts were amplified from a scaffold-tRNA$_{gly}$ template plasmid (pCD4) using primers with external Bsa1 restriction sites and purified. The backbone (pCD1) was pre-digested with Bsa1 to remove the dropout and combined with parts for Golden Gate assembly. Pre-cloned plasmids were linearized with AatII and SbfI prior to CRAPS transformations [2].

**Table S1 – Sanger-sequencing results of CRAPS variant integration**

| Series | Sample | T8 | T6 | T9 | T10 | Possible Products |
|---|---|---|---|---|---|---|
| 0 | 1 | crtI$_{Xd}$ | | | | lycopene |
| | 2 | crtI$_{Xd}$ | | | | lycopene |
| | 3 | crtI$_{Xd}$ | | | | lycopene |
| | 4 | crtI14 | | | | lycopene/2, 2, 4, -didehydrolycopene/tetradehydrolycopene |
| 1 | 1 | crtI$_{Xd}$ | crtY2 | crtZ$_{Pa}$ | tHMG1$_{Sc}$ | lycopene → β-carotene → zeaxanthin/β-cryptoxanthin |
| | 2 | crtI$_{Xd}$ | crtYB$_{Xd}$ | crtU$_{Ba}$ | crtZ$_{Hp}$ | lycopene → β-carotene → isorenieratene/renierapurpurin + zeaxanthin |
| | 3 | crtI$_{Xd}$ | crtYB$_{Xd}$ | crtU$_{Ba}$ | crtX$_{Pa}$ | lycopene → β-carotene → isorenieratene/renierapurpurin |
| | 4 | crtI$_{Xd}$ | crtY$_{Pa}$ | crtO$_{Ss}$ | crtX$_{Pa}$ | lycopene → β-carotene → canthaxanthin/echinone |
| 2 | 1 | crtI$_{Xd}$ | crtY2 | Bkt$_{Hp}$ | | lycopene → β-carotene → canthaxanthin |
| | 2 | crtI$_{Xd}$ | crtYB$_{Xd}$ | crtU$_{Ba}$ | | lycopene → β-carotene → isorenieratene/renierapurpurin |
| | 3 | crtI$_{Xd}$ | crtY$_{Pa}$ | Bkt$_{Hp}$ | | lycopene → β-carotene → canthaxanthin |
| | 4 | crtI14 | crtY2 | crtO$_{Ss}$ | | lycopene/2, 2, 4, -didehydrolycopene/tetradehydrolycopene → β-carotene → canthaxanthin/echinone |
| 3 | 1 | crtI$_{Xd}$ | crtYB$_{Xd}$ | | | lycopene → β-carotene |

| | | | | | | |
|---|---|---|---|---|---|---|
| | 2 | crtI$_{Pa}$ | crtY$_{Pa}$ | [redacted] | [redacted] | lycopene → β-carotene |
| | 3 | crtI$_{Xd}$ | crtY$_{Pa}$ | [redacted] | [redacted] | lycopene → β-carotene |
| | 4 | crtI14 | crtY2 | [redacted] | [redacted] | lycopene/2, 2, 4, -didehydrolycopene/tetradehydrolycopene → β-carotene |
| 4 | 1 | crtI$_{Pa}$ | [redacted] | [redacted] | crtI$_{Xd}$ | lycopene |
| | 2 | crtI$_{Xd}$ | [redacted] | [redacted] | tHMG1$_{Sc}$ | lycopene |
| | 3 | crtI14 | [redacted] | [redacted] | tHMG1$_{Sc}$ | lycopene/2, 2, 4, -didehydrolycopene/tetradehydrolycopene |
| | 4 | crtI$_{Xd}$ | [redacted] | [redacted] | crtE03M | lycopene |
| 5 | 1 | crtI$_{Xd}$ | crtY2 | [redacted] | crtI$_{Xd}$ | lycopene → β-carotene |
| | 2 | crtI$_{Xd}$ | crtYB$_{Xd}$ | [redacted] | crtX$_{Pa}$ | lycopene → β-carotene |
| | 3 | crtI$_{Xd}$ | crtYB$_{Xd}$ | [redacted] | tHMG1$_{Sc}$ | lycopene → β-carotene |
| | 4 | crtI14 | crtYB$_{Xd}$ | [redacted] | BTS1$_{Sc}$ | lycopene/2, 2, 4, -didehydrolycopene/tetradehydrolycopene → β-carotene |

**Table S2 – Plasmids utilized in this study**

| Plasmid | Description | Source or reference |
|---|---|---|
| pCAS-G418 | $P_{RNR2}$-*cas9*$_{NLS}$-$T_{CYC1}$, pUC, 2µ, $P_{tRNA\_Tyr}$-3′HDV-gRNA-Scaffold-$T_{SNR52}$, $P_{TEF1}$-*kanMX*-$T_{TEF1}$ | [3] |
| pCAS-Hyg | $P_{RNR2}$-*cas9*$_{NLS}$-$T_{CYC1}$, pUC, 2µ, $P_{tRNA\_Tyr}$-3′HDV-gRNA-Scaffold-$T_{SNR52}$, $P_{TEF1}$-*HphNTI*-$T_{TEF1}$ | [4] |
| pCas-split-G418 | $P_{RNR2}$-*cas9*$_{NLS}$-$T_{CYC1}$, pUC, 2µ, $P_{tRNA\_Tyr}$-3′HDV-gRNA-Scaffold-$T_{SNR52}$, $P_{TEF1}$-Split-*kanMX*-$T_{TEF1}$, $Amp^R$ | [2] |
| pCD1 | $P_{RNR2}$-*cas9*$_{NLS}$-$T_{CYC1}$, pUC, 2µ, $P_{SNR52}$-gRNA-Scaffold-$T_{SNR52}$, $P_{TEF1}$-Split-*kanMX*-$T_{TEF1}$, $Amp^R$ | This study |
| pCD3 | gRNA8 in pCas-split-G418 | This study |
| pCD4 | Guide-RNA scaffold and $tRNA_{gly}$ in pJET1.2 | [1] |
| pCD5 | gRNA8+gRNA6+gRNA9+gRNA10 in pCas-split-G418 | This study |
| pCD6 | gRNA8+gRNA6+gRNA9 in pCas-split-G418 | This study |
| pCD7 | gRNA8+gRNA6 in pCas-split-G418 | This study |
| pCD8 | gRNA8+gRNA10 in pCas-split-G418 | This study |
| pCD9 | gRNA8+gRNA6+gRNA10 in pCas-split-G418 | This study |

**Table S3 – Strains utilized in this study**

| ID | Description | Parent | Relevant genotype[a] | Reference |
|---|---|---|---|---|
| CEN.PK2-1D | Quadruple auxotroph | - | MATα his3Δ1 leu2-3_112 trp1-289 ura3-52 MAL2-8c SUC2 | |
| LP18 | CR host strain | CEN.PK2-1D | FgF20-LP1.T9-FgF20 | This study |
| LP59 | CR test strain | LP18 | FgF24-LP1.REN-FgF24 | This study |
| LP7 | 2· CR strain | LP59 | FgF18-LP1.REN-FgF18 | This study |
| LP8 | 4· CR strain | LP7 | FgF7-LP1.REN-FgF7<br>FgF19-LP1.REN-FgF19 | This study |
| LP22 | 9· CR strain | LP8 | FgF12-LP1.REN-FgF12<br>FgF16-LP1.REN-FgF16<br>FgF21-LP1.REN-FgF21<br>FgF22-LP1.REN-FgF22<br>FgF23-LP1.REN-FgF23 | This study |
| LP63 | 60 bp homology CR strain | LP18 | FgF7-60bpLP1.REN-FgF7 | This study |
| LP64 | 90 bp homology CR strain | LP18 | FgF7-90bpLP1.REN-FgF7 | This study |
| LP65 | 150 bp homology CR strain | LP18 | FgF7-150bpLP1.REN-FgF7 | This study |
| LP66 | 280 bp homology CR strain | LP18 | FgF7-280bpLP1.REN-FgF7 | This study |
| LP62 | Intermediate CRAPS strain | CEN.PK2-1D | FgF19-P$_{TDH3}$-T$_{SYNTH24}$-T10-T$_{TPS1}$-FgF19<br>FgF20-P$_{FBA1}$-T$_{FUM1}$-T8-T$_{PRM9}$-FgF20 | This study |
| LP71 | CRAPS strain | LP62 | FgF18-P$_{PDC1}$-T$_{SYNTH13}$-T9-T$_{GRE3}$-FgF18<br>FgF24-P$_{PGK1}$-T$_{SYNTH11}$-T6-T$_{HSP26}$-FgF24 | This study |
| LP75 | CRAPS *GFP* repair strain | LP71 | FgF7-P$_{FBA1}$-*GFP*-T$_{PRM9}$-FgF7 | This study |
| LP76 | CRAPS *GFP* repair strain | LP71 | FgF7-P$_{PGK1}$-*GFP*-T$_{HSP26}$-FgF7 | This study |
| LP77 | CRAPS *GFP* repair strain | LP71 | FgF7-P$_{PDC1}$-*GFP*-T$_{GRE3}$-FgF7 | This study |
| LP78 | CRAPS *GFP* repair strain | LP71 | FgF7-P$_{TDH3}$-*GFP*-T$_{TPS1}$-FgF7 | This study |
| LP86 | *crtB*+*crtE* CRAPS host | LP71 | USERXII-1-P$_{TEF1}$-*crtB*$_{Pa}$-T$_{CYC1}$-P$_{TDH3}$-*crtE*$_{Xd}$-T$_{CYC1}$-USERXII-1 | This study |
| LP191 | CRAPS carotenoid gene string 3 | LP86 | USERXII-2-P$_{PGK1}$-crtA$_{Rc}$-T$_{HSP26}$-P$_{PDC1}$-crtZ$_{Pa}$-T$_{GRE3}$-P$_{TDH3}$-crtI$_{Xd}$-T$_{TPS1}$-USERXII-2 | This study |

| | | | | |
|---|---|---|---|---|
| LP222 | CRAPS carotenoid gene string 1 | LP191 | FgF21-$\underline{P_{FBA1}}$-crt$I_{Xd}$-$\underline{T_{PRM9}}$-$\underline{P_{PGK1}}$-crt$YB_{Xd}$-$\underline{T_{HSP26}}$-$\underline{P_{PDC1}}$-crt$U_{Ba}$-$\underline{T_{GRE3}}$-$\underline{P_{TDH3}}$-tHMG$1_{Sc}$-$\underline{T_{TPS1}}$-FgF21 | This study |
| LP309 | CRAPS carotenoid gene string 4 | LP222 | FgF16-$\underline{P_{FBA1}}$-crt$I_{14}$-$\underline{T_{PRM9}}$-$\underline{P_{PGK1}}$-crtY2-$\underline{T_{HSP26}}$-$\underline{P_{PDC1}}$-crtE03M-$\underline{T_{GRE3}}$-$\underline{P_{TDH3}}$-BTS$1_{Sc}$-$\underline{T_{TPS1}}$-FgF16 | This study |
| LP368 | CRAPS carotenoid gene string 2 | LP309 | FgF7-$\underline{P_{FBA1}}$-crt$I_{Pa}$-$\underline{T_{PRM9}}$-$\underline{P_{PGK1}}$-crt$Y_{Pa}$-$\underline{T_{HSP26}}$-$\underline{P_{PDC1}}$-crt$O_{Ss}$-$\underline{T_{GRE3}}$-$\underline{P_{TDH3}}$-crt$X_{Pa}$-$\underline{T_{TPS1}}$-FgF7 | This study |
| LP508 | CRAPS carotenoid gene string 5 | LP368 | USERXII-5-$\underline{P_{FBA1}}$-crt$I_{Nc}$-$\underline{T_{PRM9}}$-$\underline{P_{PGK1}}$-crtYB01M-$\underline{T_{HSP26}}$-$\underline{P_{PDC1}}$-tHMG$1_{Sc}$-$\underline{T_{GRE3}}$-$\underline{P_{TDH3}}$-crtI14-$\underline{T_{TPS1}}$-USERXII-5 | This study |
| LP526 | CRAPS carotenoid gene string 7 | LP508 | 511b-$\underline{P_{PGK1}}$-crtOx-$\underline{T_{HSP26}}$-$\underline{P_{TDH3}}$-crtE03M-$\underline{T_{TPS1}}$-511b | This study |
| LP556 | CRAPS carotenoid gene string 6 | LP526 | 911b-$\underline{P_{PDC1}}$-BKT$_{Hp}$*-$\underline{T_{GRE3}}$-$\underline{P_{TDH3}}$-crt$Z_{Hp}$-$\underline{T_{TPS1}}$-911b | This study |

[a] Truncated CRAPS promoter and terminator elements are underlined.

**Table S4 – *S. cerevisiae* integration sites utilized in this study**

| Target site ID | Target site sequence[a] | Reference |
|---|---|---|
| FgF7 | TATCCTGAATGTTCTCTCCCAGG | [5, 6] |
| FgF12 | TGTCTTGTTAGAATGAAGATCGG | [5] |
| FgF16 | TGTACCAAAAGTTATCCTGTAGG | [5, 6] |
| FgF18 | ATAGAATTACTATTGAAGAGTGG | [5, 6] |
| FgF19 | ATTCACTCTGCTAAGATTATCGG | [5, 6] |
| FgF20 | GTTAGAGCTGTTACAAGTTACGG | [5, 6] |
| FgF21 | TCTTGGGTTGCCAAACTAAGAGG | [5] |
| FgF22 | GGGTCAACAGTATAGAACCGTGG | [5] |
| FgF23 | TATTTTCAATATTATACAAGGG | [5] |
| FgF24 (*PDC6*) | GTACAACGAAATCCAGACCTGGG | [5, 6] |
| USERXII-1 | GTCTTTGCCGGTTACCCATCTGG | [7] |
| USERXII-2 | TCGAGAGAGTCGCCGATAGTAGG | [7] |
| USERXII-5 | TTGTCACAGTGTCACATCAGCGG | [7] |
| 511b | CAGTGTATGCCAGTCAGCCACGG | [8] |
| 911b | GTAATATTGTCTTGTTTCCCTGG | [8] |

[a] Protospacer-adjacent motifs are underlined.

**Table S5 – Genes and synthetic DNAs synthesized in this study**

| DNA | Source | Ref | Codon opt. | Sequence (5′ – 3′)[a] |
|---|---|---|---|---|
| $crtB_{Pa}$ | *Pantoea ananatis* | [9] | Yes | ATGAATAATCCGTCTCTTTTAAATCATGCTGTGGAAACGATGGCTGTTG GGAGTAAATCTTTCGCAACGGCCTCAAAATTGTTTGACGCGAAAACAA GGAGGAGCGTTCTAATGCTGTACGCCTGGTGTCGTCATTGTGATGACG TAATCGACGACCAAACCTTAGGGTTTCAGGCCAGACAGCCGGCGTTGC AGACCCCCGAACAAAGACTAATGCAGTTAGAGATGAAGACTAGACAA GCGTATGCCGGGAGCCAGATGCACGAACCGGCCTTCGCTGCGTTTCAG GAAGTCGCGATGGCCCACGACATCGCACCCGCCTATGCGTTTGACCAT TTAGAGGGTTTTGCAATGGATGTTAGGGAGGCTCAATATAGTCAATTG GACGATACCCTTAGATACTGCTATCATGTGGCAGGAGTTGTGGGGCTG ATGATGGCCCAGATCATGGGTGTTAGAGACAATGCGACCCTTGATAGG GCCTGCGATCTGGGCTTAGCTTTCCAGCTGACGAACATAGCGAGAGAC ATCGTTGATGATGCTCACGCAGGGCGTTGCTACTTACCTGCGTCCTGGC TAGAGCACGAGGGCCTGAATAAGGAAAACTACGCGGCTCCCGAGAAT CGTCAAGCGCTTTCCAGAATAGCGAGAAGACTGGTTCAGGAGGCAGA ACCTTATTACCTTTCTGCAACAGCGGGGTTAGCGGGACTGCCCCTTAG ATCTGCATGGGCTATCGCGACTGCTAAACAGGTCTATAGAAAAATCGG AGTCAAGGTCGAACAGGCGGGGCAACAAGCGTGGGATCAGAGACAGT CCACCACCACTCCAGAAAAATTGACCCTGCTTTTAGCAGCTTCTGGGC AGGCATTAACGTCAAGAATGAGGGCTCACCCACCACGTCCGGCTCACT TGTGGCAGAGGCCCTTGGGCTGA |
| $crtE_{Xd}$ | *Xanthophyllomyces dendrorhous* | [10] | Yes | ATGGATTACGCGAACATCCTCACAGCAATTCCACTCGAGTTTACTCCTC AGGATGATATCGTGCTCCTTGAACCGTATCACTACCTAGGAAAGAACC CTGGAAAAGAAATTCGATCACAACTCATCGAGGCTTTCAACTATTGGT TGGATGTCAAGAAGGAGGATCTCGAGGTCATCCAGAACGTTGTTGGCA TGCTACATACCGCTAGCTTATTAATGGACGATGTGGAGGATTCATCGG TCCTCAGGCGTGGGTCGCCTGTGGCCCATCTAATTTACGGGATTCCGCA GACAATAAACACTGCAAACTACGTCTACTTTCTGGCTTATCAAGAGAT |

| | | | | |
|---|---|---|---|---|
| | | | | CTTCAAGCTTCGCCCAACACCGATACCCATGCCTGTAATTCCTCCTTCA<br>TCTGCTTCGCTTCAATCATCCGTCTCCTCTGCATCCTCCTCCTCCTCGGC<br>CTCGTCTGAAAACGGGGGCACGTCAACTCCTAATTCGCAGATTCCGTT<br>CTCGAAAGATACGTATCTTGATAAAGTGATCACAGACGAGATGCTTTC<br>CCTCCATAGAGGGCAGGGCCTGGAGCTATTCTGGAGAGATAGTCTGAC<br>GTGTCCTAGCGAAGAGGAATATGTGAAAATGGTTCTTGGAAAGACGG<br>GAGGTTTGTTCCGTATAGCGGTCAGATTGATGATGGCAAAGTCAGAAT<br>GTGACATAGACTTTGTCCAGCTTGTCAACTTGATCTCAATATACTTCCA<br>GATCAGGGATGACTATATGAACCTTCAGTCTTCTGAGTATGCCCATAA<br>TAAGAATTTTGCAGAGGACCTCACAGAAGGGAAATTCAGTTTTCCCAC<br>TATCCACTCGATTCATGCCAACCCCTCATCGAGACTCGTCATCAATACG<br>TTGCAGAAGAAATCGACCTCTCCTGAGATCCTTCACCACTGTGTAAAC<br>TACATGCGCACAGAAACCCACTCATTCGAATATACTCAGGAAGTCCTC<br>AACACCTTGTCAGGTGCACTCGAGAGAGAACTAGGAAGGCTTCAAGG<br>AGAGTTCGCAGAAGCTAACTCAAGGATGGATCTTGGAGACGTAGATTC<br>GGAAGGAAGAACGGGGAAGAACGTCAAATTGGAAGCGATCCTGAAAA<br>AGCTAGCCGATATCCCTCTGTGA |
| crtI$_{Xd}$ | *Xanthophyllomyces dendrorhous* | [10] | Yes | ATGGGAAAAGAACAAGATCAGGATAAACCCACAGCTATCATCGTGGG<br>ATGTGGTATCGGTGGAATCGCCACTGCCGCTCGTCTTGCTAAAGAAGG<br>TTTCCAGGTCACGGTGTTCGAGAAGAACGACTACTCCGGAGGTCGATG<br>CTCTTTAATCGAGCGAGATGGTTATCGATTCGATCAGGGGCCCAGTTT<br>GCTGCTCTTGCCAGATCTCTTCAAGCAGACATTCGAAGATTTGGGAGA<br>GAAGATGGAAGATTGGGTCGATCTCATCAAGTGTGAACCCAACTATGT<br>TTGCCACTTCCACGATGAAGAGACTTTCACTTTTTCAACCGACATGGCG<br>TTGCTCAAGCGGGAAGTCGAGCGTTTTGAAGGCAAAGATGGATTTGAT<br>CGGTTCTTGTCGTTTATCCAAGAAGCCCACAGACATTACGAGCTTGCTG<br>TCGTTCACGTCCTGCAGAAGAACTTCCCTGGCTTCGCAGCATTCTTACG<br>GCTACAGTTCATTGGCCAAATCCTGGCTCTTCACCCCTTCGAGTCTATC<br>TGGACAAGAGTTTGTCGATATTTCAAGACCGACAGATTACGAAGAGTC<br>TTCTCGTTTGCAGTGATGTACATGGGTCAAAGCCCATACAGTGCGCCC<br>GGAACATATTCCTTGCTCCAATACACCGAATTGACCGAGGGCATCTGG<br>TATCCGAGAGGAGGCTTTTGGCAGGTTCCTAATACTCTTCTTCAGATCG<br>TCAAGCGCAACAATCCCTCAGCCAAGTTCAATTTCAACGCTCCAGTTTC |

| | | | | |
|---|---|---|---|---|
| | | | | CCAGGTTCTTCTCTCTCCTGCCAAGGACCGAGCGACTGGTGTTCGACTT<br>GAATCCGGCGAGGAACATCACGCCGATGTTGTGATTGTCAATGCTGAC<br>CTCGTTTACGCCTCCGAGCACTTGATTCCTGACGATGCCAGAAACAAG<br>ATTGGCCAACTGGGTGAAGTCAAGAGAAGTTGGTGGGCTGACTTAGTT<br>GGTGGAAAGAAGCTCAAGGGAAGTTGCAGTAGTTTGAGCTTCTACTGG<br>AGCATGGACCGAATCGTGGACGGTCTGGGCGGACACAATATCTTCTTG<br>GCCGAGGACTTCAAGGGATCATTCGACACAATCTTCGAGGAGTTGGGT<br>CTCCCAGCCGATCCTTCCTTTTACGTGAACGTTCCCTCGCGAATCGATC<br>CTTCTGCCGCTCCCGAAGGCAAAGATGCTATCGTCATTCTTGTGCCGTG<br>TGGCCATATCGACGCTTCGAACCCTCAAGATTACAACAAGCTTGTTGC<br>TCGGGCAAGGAAGTTTGTGATCCAAACGCTTTCCGCCAAGCTTGGACT<br>TCCCGACTTTGAAAAAATGATTGTGGCAGAGAAGGTTCACGATGCTCC<br>CTCTTGGGAGAAAGAATTTAACCTCAAGGACGGAAGCATCTTGGGACT<br>GGCTCACAACTTTATGCAAGTTCTTGGTTTCAGGCCGAGCACCAGACA<br>TCCCAAGTATGACAAGTTGTTCTTTGTCGGGGCTTCGACTCATCCCGGA<br>ACTGGGGTTCCCATCGTCTTGGCTGGAGCCAAGTTAACTGCCAACCAA<br>GTTCTCGAATCCTTTGACCGATCCCCAGCTCCAGATCCCAATATGTCAC<br>TCTCCGTACCATATGGAAAACCTCTCAAATCAAATGGAACGGGTATCG<br>ATTCTCAGGTCCAGCTGAAGTTCATGGATTTGGAGAGATGGGTATACC<br>TTTTGGTGTTGTTGATTGGGGCCGTGATCGCTCGATCCGTTGGTGTTCT<br>TGCTTTCTGA |
| crtYB$_{Xd}$ | *Xanthophyllo myces dendrorhous* | [11] | Yes | ATGACGGCTCTCGCATATTACCAGATCCATCTGATCTATACTCTCCCAA<br>TTCTTGGTCTTCTCGGCCTGCTCACTTCCCCGATTTTGACAAAATTGA<br>CATCTACAAAATATCGATCCTCGTATTTATTGCGTTTAGTGCAACCACA<br>CCATGGGACTCATGGATCATCAGAAATGGCGCATGGACATATCCATCA<br>GCGGAGAGTGGCCAAGGCGTGTTTGGAACGTTTCTAGATGTTCCATAT<br>GAAGAGTACGCTTTCTTTGTCATTCAAACCGTAATCACCGGCTTGGTCT<br>ACGTCTTGGCAACTAGGCACCTTCTCCCATCTCTCGCGCTTCCCAAGAC<br>TAGATCGTCCGCCCTTTCTCTCGCGCTCAAGGCGCTCATCCCTCTGCCC<br>ATTATCTACCTATTTACCGCTCACCCCAGCCCATCGCCCGACCCGCTCG<br>TGACAGATCACTACTTCTACATGCGGGCACTCTCCTTACTCATCACCCC<br>ACCTACCATGCTCTTGGCAGCATTATCAGGCGAATATGCTTTCGATTGG<br>AAAAGTGGCCGAGCAAAGTCAACTATTGCAGCAATCATGATCCCGACG |

| | | | | |
|---|---|---|---|---|
| | | | | GTGTATCTGATTTGGGTAGATTATGTTGCTGTCGGTCAAGACTCTTGGT<br>CGATCAACGATGAGAAGATTGTAGGGTGGAGGCTTGGAGGTGTACTAC<br>CCATTGAGGAAGCTATGTTCTTCTTACTGACGAATCTAATGATTGTTCT<br>GGGTCTGTCTGCCTGCGATCATACTCAGGCCCTATACCTGCTACACGGT<br>CGAACTATTTATGGCAACAAAAAGATGCCATCTTCATTTCCCCTCATTA<br>CACCGCCTGTGCTCTCCCTGTTTTTAGCAGCCGACCATACTCTTCTCA<br>GCCAAAACGTGACTTGGAACTGGCAGTCAAGTTGTTGGAGAAAAAGA<br>GCCGGAGCTTTTTTGTTGCCTCGGCTGGATTTCCTAGCGAAGTTAGGGA<br>GAGGCTGGTTGGACTATACGCATTCTGCCGGGTGACTGATGATCTTAT<br>CGACTCTCCTGAAGTATCTTCCAACCCGCATGCCACAATTGACATGGTC<br>TCCGATTTTCTTACCCTACTATTTGGGCCCCCGCTACACCCTTCGCAAC<br>CTGACAAGATCCTTTCTTCGCCTTTACTTCCTCCTTCGCACCCTTCCCGA<br>CCCACGGGAATGTATCCCCTCCCGCCTCCTCCTTCGCTCTCGCCTGCCG<br>AGCTCGTTCAATTCCTTACCGAAAGGGTTCCCGTTCAATACCATTTCGC<br>CTTCAGGTTGCTCGCTAAGTTGCAAGGGCTGATCCCTCGATACCCACTC<br>GACGAACTCCTTAGAGGATACACCACTGATCTTATCTTTCCCTTATCGA<br>CAGAGGCAGTCCAGGCTCGGAAGACGCCTATCGAGACCACAGCTGAC<br>TTGCTGGACTATGGTCTATGTGTAGCAGGCTCAGTCGCCGAGCTATTG<br>GTCTATGTCTCTTGGGCAAGTGCACCAAGTCAGGTCCCTGCCACCATA<br>GAAGAAAGAGAAGCTGTGTTAGTGGCAAGCCGAGAGATGGGAACTGC<br>CCTTCAGTTGGTGAACATTGCTAGGGACATTAAAGGGGACGCAACAGA<br>AGGGAGATTTTACCTACCACTCTCATTCTTTGGTCTTCGGGATGAATCA<br>AAGCTTGCGATCCCGACTGATTGGACGGAACCTCGGCCTCAAGATTTC<br>GACAAACTCCTCAGTCTATCTCCTTCGTCCACATTACCATCTTCAAACG<br>CCTCAGAAAGCTTCCGGTTCGAATGGAAGACGTACTCGCTTCCATTAG<br>TCGCCTACGCAGAGGATCTTGCCAAACATTCTTATAAGGGAATTGACC<br>GACTTCCTACCGAGGTTCAAGCGGGAATGCGAGCGGCTTGCGCGAGCT<br>ACCTACTGATCGGCCGAGAGATCAAAGTCGTTTGGAAAGGAGACGTCG<br>GAGAGAGAAGGACAGTTGCCGGATGGAGGAGAGTACGGAAAGTCTTG<br>AGTGTGGTCATGAGCGGATGGGAAGGGCAGTAA |
| crtU$_{Ba}$ | *Brevibacterium aurantiacum* | [12] | Yes | ATGACACAAAGGAGAAGACCCAGGGATAGATTCGCCGAAAGAATTCA<br>AGGACCTCAAGGTAGACCGAGGCTTTTAAGACCAAAGAGGGTCACAA<br>TCATAGGTGCAGGAATAGCGGGCTTGGCTGCCGCAGCTATTCTTGCCG |

| | | | | |
|---|---|---|---|---|
| | | | | AACACGGGGCCGAGGTAACCGTAATTGAGAAGACTGATTACCTTGGTG<br>GGCGTGTTGGTGCGTGGCCGGTGGATGACGAACGTACGATGTCCAGGG<br>GCTTCCACGCTTTTTTCAGGCAATATTACAATCTTCGTGACCTATTGTC<br>AAGGGCCGATCCGGAAGGGGAGTGTCTAAGACCAGTCGACGACTATC<br>CTCTGATACACAGAAGGGGCTCTATGGATACTTTTGCATCTATTCCCAG<br>GACGCCTCCTTTTAATCTTTTAGGCTTCGTGTGGCAGTCACCCACTTTC<br>CCGATTCGTGGACTTAGAGACGTAGATATCGCGGCAGCCGTCGAGCTG<br>ATAGACGTAGAATTCCCTGCTACATATTCTTATTACGATGGCGAATCCG<br>CGGCAGATTTCTTGGATCGTCTGAGGTTCCCCGATGAGGCGAGACACC<br>TGGCCTTGGAGGTTTTCGCTAGGTCTTTCTTTGCCGACCCAACAGAGTT<br>CTCCGCAGGAGAATTGGTAGCAATGTTCCACACGTACTTCACCGGTTC<br>TGCCGAGGGACTATTATTTGATGTCCCGGTAGACGACTATGACACCGC<br>ACTGTGGGCACCCCTGGGCGGGTATCTTGAGTCACTGGGAGTGACAAT<br>TGAAACCGGAACAACGGTCACATCTATTGACCCAACCGAATCCGGGTG<br>GACTACCACGACAGGAGAGGCTAATTTGGAAAGCGACGCGGTTGTGCT<br>GGCAGTAGACCCTGCCGCAGCAAGGGATTTGTTAAGCGCATCACATGA<br>TAGCCTTGTAGACTCTGCTCCGGCAGCACAAAGATGGATGGAGACTAT<br>TGGTTCACAAACAAATGCACCAGCGTTTGCAGTTTTGAGATTATGGCTT<br>GGAACCCCCGTCGCAGATCACAGACCGGCCTTTCTAGGTACGTCTGGA<br>TACGATCTGTTGGATAACGTGAGTGTACTTGAGCGTTTCGAGGCTGGT<br>GCAAGGGCTTGGTCTGAGTCCCATCATGGGAGTGTGTTAGAGCTGCAT<br>GCATACGCTCTGGAAGGCGACAGTTACGATACCGAGAGGGGTAGAGC<br>TGATATCGTTGCCAGGTTACTATCTGACCTGCACCATGTGTACCCTGAA<br>ACGGCAGCATTAACCATTGTGGACCAGGAACTGCTGATTGAGGCCGAT<br>TGCGGCCTAACAGATACTCGTCCGTGGGAAGATAGGCCGGAGCCGTCT<br>ACGCCCATCCCAGGCCTGGTAGTTGCGGGAGACTATGTCAGGTGCAAC<br>ACTCCCGTCGCATTAATGGAGCGTGCAGCCACTACAGGGTATCTGGCG<br>GCAAACCATCTATTGTCCACCTGGCGTGTCGAAGGTACAGACCTATGG<br>TCACCTCCTACAAGGGGCCTTCTGAGACGTGGCGTTCTTGGTTTAATAA<br>GGCGTCGTAGGTAA |
| crtI$_{Pa}$ | *Pantoea ananatis* | [9] | Yes | ATGAAGCCCACGACGGTAATAGGCGCTGGTTTTGGAGGATTAGCTTTG<br>GCCATTCGTTTGCAGGCCGCTGGCATTCCCGTCCTATTGTTAGAACAGA<br>GGGACAAGCCCGGTGGTCGTGCCTACGTCTATGAAGACCAGGGCTTCA |

| | | | | |
|---|---|---|---|---|
| | | | | CATTCGACGCAGGACCAACGGTAATAACTGACCCTTCAGCTATAGAGG<br>AGTTATTCGCCCTTGCCGGGAAGCAATTAAAAGAATATGTTGAGCTAC<br>TGCCGGTTACACCCTTCTATAGGCTTTGTTGGGAGTCTGGCAAAGTGTT<br>CAACTATGATAACGATCAGACCCGTCTGGAAGCGCAAATCCAGCAGTT<br>TAATCCCAGGGACGTTGAAGGTTACAGACAGTTCCTTGATTACAGTAG<br>AGCTGTCTTTAAGGAAGGATACTTGAAATTAGGGACGGTCCCCTTCCT<br>ATCCTTCCGTGATATGCTGCGTGCAGCTCCTCAGTTAGCTAAACTGCAA<br>GCATGGCGTTCCGTTTACTCTAAGGTAGCATCCTACATTGAAGATGAG<br>CACTTGAGACAGGCGTTCTCTTTTCACTCTCTTCTGGTAGGCGGGAATC<br>CTTTCGCAACCAGTTCCATATACACACTTATTCACGCACTGGAACGTGA<br>GTGGGGAGTTTGGTTTCCTCGTGGGGGCACGGGTGCGTTGGTCCAAGG<br>AATGATTAAGCTGTTCCAGGATCTAGGAGGTGAAGTCGTCCTTAACGC<br>GCGTGTGTCACATATGGAAACTACTGGAAACAAAATAGAAGCGGTCC<br>ATTTGGAAGATGGCAGGAGATTCTTAACTCAGGCAGTCGCTTCCAATG<br>CGGATGTTGTTCATACTTACAGGGACTTACTTTCCCAACACCCCGCTGC<br>CGTAAAGCAATCAAACAAACTACAGACGAAGAGAATGTCTAACAGCC<br>TGTTTGTTTTATATTTCGGATTGAATCACCATCATGACCAGCTTGCACA<br>TCATACCGTATGTTTTGGTCCAAGATACAGGGAATTAATCGATGAGAT<br>ATTCAATCACGACGGACTTGCTGAAGACTTTAGTTTATACCTTCATGCT<br>CCCTGCGTGACCGATAGTAGTTTAGCGCCAGAGGGTTGTGGGTCATAT<br>TACGTGTTGGCTCCGGTACCACACCTAGGAACAGCAAATCTGGACTGG<br>ACAGTGGAGGGTCCTAAACTACGTGACCGTATCTTTGCTTATTTAGAA<br>CAGCATTATATGCCCGGGTTACGTAGTCAATTGGTGACACATCGTATG<br>TTTACGCCCTTTGATTTTCGTGACCAATTGAATGCTTACCACGGCTCCG<br>CTTTTTCCGTTGAGCCGGTTCTAACACAGTCCGCATGGTTCAGGCCCCA<br>CAATCGTGATAAAACGATCACCAACCTTTATCTAGTAGGTGCAGGCAC<br>ACACCCGGGTGCGGGCATACCCGGCGTTATCGGCAGTGCAAAGGCGA<br>CCGCGGGTCTTATGCTAGAAGATCTAATCGGCTGA |
| $crtY_{Pa}$ | Pantoea ananatis | [13] | Yes | ATGCAACCGCATTATGACTTGATACTGGTTGGAGCAGGACTTGCGAAT<br>GGGTTGATTGCACTACGTCTACAGCAACAACAGCCAGATATGAGAATA<br>CTTCTAATAGATGCAGCTCCACAGGCGGGCGGTAACCATACGTGGTCC<br>TTCCACCACGATGATTTGACAGAAAGCCAACACAGATGGATCGCACCC<br>CTGGTAGTACATCATTGGCCGGACTACCAGGTACGTTTCCCCACAAGG |

| | | | | |
|---|---|---|---|---|
| | | | | AGGAGAAAACTAAACTCAGGTTACTTTTGCATAACTTCCCAAAGATTC GCAGAGGTGCTACAACGTCAATTCGGTCCGCATCTTTGGATGGATACT GCGGTTGCCGAAGTGAACGCCGAGTCCGTGCGTCTTAAGAAGGGTCAG GTCATCGGCGCGCGTGCTGTTATCGACGGCAGGGGGTATGCAGCTAAC TCTGCATTATCAGTCGGGTTTCAAGCGTTTATCGGACAGGAGTGGAGA CTTTCCCACCCGCACGGCTTAAGCAGCCCAATAATAATGGACGCCACT GTGGATCAGCAAAATGGGTATAGATTCGTTTACAGTTTGCCCCTGTCTC CCACTAGACTACTGATTGAGGATACTCATTATATTGATAACGCGACTCT AGACCCCGAGTGTGCCCGTCAGAATATTTGTGACTACGCCGCGCAACA AGGCTGGCAACTTCAAACTCTATTAAGAGAAGAACAGGGTGCGCTGCC AATCACTTTATCTGGTAACGCTGATGCATTTTGGCAGCAACGTCCCCTT GCGTGTAGTGGTCTTAGGGCAGGGTTGTTTCACCCCACGACTGGTTATT CATTACCCTTAGCCGTTGCGGTAGCCGACAGACTTTCCGCCCTGGATGT CTTTACCTCTGCTAGCATACATCACGCAATCACGCATTTCGCAAGAGA AAGATGGCAGCAACAAGGTTTCTTTAGAATGCTTAATCGTATGCTATTT CTAGCTGGACCTGCTGATTCACGTTGGCGTGTTATGCAAAGGTTCTATG GTTTACCAGAGGATTTAATTGCACGTTTCTATGCAGGCAAACTGACTTT AACCGACAGGTTGCGTATTTTATCCGGGAAGCCCCCGGTCCCAGTACT TGCAGCTCTACAAGCGATTATGACTACTCATAGAGGCTGA |
| crtO$_{Ss}$ | *Synechocystis sp.* (unclassified) | [14] | Yes | ATGATAACTACAGACGTGGTTATTATCGGCGCGGGGCATAACGGACTT GTGTGTGCTGCGTATCTACTTCAGCGTGGCTTGGGAGTAACATTGCTGG AAAAGAGAGAGGTCCCTGGTGGAGCCGCAACAACAGAGGCACTGATG CCAGAACTGAGTCCACAGTTTCGTTTCAATAGATGCGCCATCGACCAT GAATTCATATTTCTTGGACCTGTATTACAGGAACTGAATTTAGCTCAGT ACGGATTAGAATACCTATTCTGTGACCCTTCAGTTTTCTGTCCGGGGCT TGACGGGCAGGCCTTCATGTCCTACAGATCACTAGAGAAAACCTGTGC GCACATTGCAACATATAGCCCGCGTGATGCAGAAAAATATCGTCAATT CGTTAATTATTGGACAGATCTGCTGAACGCAGTCCAACCAGCATTTAA CGCGCCTCCTCAGGCCCTACTAGATTTGGCGCTAAATTATGGATGGGA AAATCTGAAGTCTGTATTGGCCATAGCAGGAAGTAAGACCAAGGCGCT TGATTTCATACGTACAATGATTGGCAGCCCAGAAGATGTCCTAAACGA ATGGTTCGACTCCGAGAGAGTTAAGGCCCCCCTAGCTCGTCTTTGTTCA GAAATCGGAGCACCGCCGAGCCAAAAGGGTTCATCCAGTGGAATGAT |

| | | | | |
|---|---|---|---|---|
| | | | | GATGGTTGCGATGCGTCATCTGGAGGGCATAGCTCGTCCGAAAGGCGG<br>TACCGGTGCTCTTACTGAAGCCCTGGTTAAGCTTGTCCAGGCCCAAGG<br>AGGCAAAATCCTTACCGACCAAACGGTGAAGCGTGTCCTTGTTGAAAA<br>CAACCAAGCGATCGGTGTCGAGGTGGCCAACGGCGAACAGTACCGTG<br>CGAAGAAGGGAGTCATTAGCAATATCGACGCACGTAGGCTATTTTTAC<br>AGCTAGTTGAGCCGGGGGCATTAGCTAAGGTGAATCAGAATTTAGGTG<br>AACGTCTGGAGAGACGTACGGTCAACAACAACGAAGCGATATTAAAA<br>ATAGACTGCGCCCTAAGTGGGTTGCCTCATTTTACCGCAATGGCCGGG<br>CCTGAAGACCTGACCGGCACAATTCTTATCGCTGATAGCGTTAGGCAC<br>GTCGAAGAGGCTCATGCGCTTATAGCGCTGGGACAGATACCAGACGCG<br>AATCCGAGCCTTTACCTAGACATCCCTACCGTGCTTGACCCCACAATG<br>GCTCCCCCTGGGCAACACACCCTATGGATTGAATTCTTTGCCCCCTATC<br>GTATTGCTGGTTTAGAGGGCACGGGCCTGATGGGGACGGGTTGGACTG<br>ACGAACTGAAGGAGAAGGTAGCAGATAGGGTCATCGATAAACTAACG<br>GACTATGCGCCCAACTTAAAGAGCTTGATAATAGGACGTCGTGTAGAA<br>TCACCGGCTGAACTTGCGCAGCGTCTAGGGAGTTACAATGGGAATGTC<br>TATCACCTGGATATGTCACTAGATCAAATGATGTTTCTTAGACCCTTAC<br>CGGAAATAGCAAACTATCAAACGCCGATTAAAAACCTATATCTAACTG<br>GAGCGGGTACGCACCCAGGGGGGTCCATCAGCGGCATGCCCGGTAGA<br>AATTGTGCTAGAGTCTTCCTAAAGCAGCAAAGGAGATTTTGGGGCTGA |
| crtX$_{Pa}$ | *Pantoea ananatis* | [15] | | ATGTCTCACTTCGCAGCTATCGCTCCCCCGTTCTATTCACACGTGCGTG<br>CCCTTCAGAATTTGGCTCAGGAGTTAGTCGCCAGGGGGCACCGTGTTA<br>CTTTCATCCAACAGTATGATATAAAGCATCTTATCGACTCAGAAACCA<br>TCGGTTTTCACTCAGTCGGCACGGATTCTCATCCGCCTGGTGCTCTGAC<br>TCGTGTTCTACACCTAGCTGCACATCCACTTGGGCCCAGTATGTTGAAA<br>TTGATCAACGAGATGGCGCGTACTACGGACATGCTTTGTAGAGAATTA<br>CCGCAGGCTTTTAACGACCTAGCTGTAGATGGTGTAATCGTCGACCAG<br>ATGGAACCTGCAGGAGCACTTGTGGCAGAGGCGCTAGGCCTTCCGTTC<br>ATTTCCGTGGCGTGCGCACTTCCTTTAAACCGTGAACCCGATATGCCCT<br>TAGCCGTCATGCCGTTCGAGTATGGTACTTCCGATGCGGCTAGGGAAA<br>GATATGCAGCAAGTGAGAAAATTTACGACTGGCTTATGCGTAGGCATG<br>ATAGGGTTATTGCAGAGCATTCCCACAGAATGGGGCTAGCCCCCAGAC<br>AAAAGCTTCATCAGTGCTTCTCCCCCTTGGCACAGATATCACAATTAGT |

| | | | | |
|---|---|---|---|---|
| | | | | ACCAGAGCTAGACTTTCCCAGGAAAGCGCTTCCTGCGTGTTTCCACGC<br>AGTTGGTCCTCTGAGAGAGACTCATGCCCCCTCAACCTCTAGTTCAAG<br>ATACTTTACATCTTCTGAAAAACCAAGGATTTTCGCTTCCCTGGGAACA<br>CTACAAGGCCATCGTTATGGCCTTTTTAAAACAATCGTAAAGGCTTGT<br>GAAGAAATTGATGGACAGCTACTGTTGGCTCACTGCGGCAGGCTGACG<br>GATAGCCAATGCGAAGAATTAGCTAGGAGTCGTCACACGCAAGTAGT<br>AGATTTCGCGGATCAGTCAGCGGCCTTGAGTCAGGCCCAGCTGGCGAT<br>CACTCACGGAGGCATGAACACGGTGCTGGATGCCATCAACTATCGTAC<br>GCCTTTGCTTGCGCTACCGTTAGCGTTTGACCAGCCCGGAGTAGCTAGT<br>CGTATTGTGTATCACGGTATCGGCAAGAGAGCTTCCCGTTTTACCACCA<br>GTCATGCACTGGCTAGACAAATGAGGTCCCTATTGACTAATGTCGATT<br>TCCAGCAGCGTATGGCCAAGATACAAACTGCGTTAAGGCTTGCTGGGG<br>GTACGATGGCAGCAGCAGACATCATTGAGCAAGTCATGTGCACAGGCC<br>AGCCGGTACTTTCTGGGTCTGGTTATGCGACCGCGTTGGGCTGA |
| $crtA_{Rc}$ | *Rhodobacter capsulatus* | [16] | Yes | ATGCCTGTAGCTTCACTATCCCTTTTCCGTTTCGACGGTACTAGTAGCT<br>TACCTTGGGTTATATCACAGATGATACTGAGCCGTAGACCCTTGAACG<br>ATGAGCCAAGAGTCAAGTTCTATAAACTTTGCGGTAGTGGGACGGGGG<br>AGGGCTTCACTCCAAAACCTAACTGGAGAGTATGGGCCATAATGGCTG<br>CCTTTGACACCGAAGCCGACGCCCGTGACGTTACGGCCAATCACCCCG<br>TTTGGAAGAGGTGGAGAGCCCACGCAGCAGAAACTCTGGTTCTTCATT<br>TACAGCCCCTATCAGCACGTGGTACGTGGGGAGGAGTTAATCCATTTT<br>TACCAGAACAGGTAGCTGAACCTAGTCCCGATGAGCCTGTAGTAGCCT<br>TGACCAGGGCGGCGATAAAACCCCACAAGGCAAATGCCTTTTGGAGC<br>AGAGTACCGAAGATAAGCGAGAAAGTCGGTGAGGACCAAAATCTTAT<br>GTTCAAGATAGGCATCGGAGAGATCCCCCTTTTCCACCAGGTTACGTTT<br>AGCATTTGGCCCGACGTGGCTAAAATGAATGCATTCGCCCGTGGAGAT<br>ACTCCACATGGTAAGGCTATTCGTGCTGCTAGAGAGGAGGGTTGGTTT<br>ACAGAAGAACTGTATGCGCGTTTCAGGTTACTGGGAACGGAGGGTAGC<br>TGGATGGGCAAGGACCCCCTTGCGTCAAAAGTACTAGAGAGAGAAAC<br>GGCGGGCTGA |
| $crtZ_{Pa}$ | *Pantoea ananatis* | [17] | Yes | ATGTTATGGATATGGAATGCTCTTATAGTATTCGTAACAGTTATTGGGA<br>TGGAAGTTATCGCAGCGTTAGCGCATAAGTACATCATGCACGGTTGGG<br>GATGGGGCTGGCACTTATCCCACCATGAACCTAGAAAAGGTGCGTTCG |

| | | | | |
|---|---|---|---|---|
| | | | | AGGTGAATGATTTGTACGCAGTAGTGTTTGCAGCTTTGTCCATACTGCT<br>TATATATCTTGGCTCAACGGGAATGTGGCCCTTACAATGGATAGGGGC<br>CGGAATGACTGCATACGGTCTGTTGTACTTCATGGTGCACGATGGCTT<br>AGTACATCAACGTTGGCCGTTTCGTTACATCCCCCGTAAAGGTTACCTG<br>AAAAGACTTTACATGGCCCATAGAATGCACCACGCCGTCAGGGGGAA<br>AGAGGGGTGCGTGTCCTTCGGCTTCCTATATGCACCCCCCTTGTCTAAG<br>CTGCAGGCAACACTTAGAGAGAGACATGGGGCCAGGGCTGGGGCAGC<br>CCGTGATGCGCAAGGAGGGGAAGACGAGCCAGCTTCTGGAAAAGGCT<br>GA |
| crtI14 | *Pantoea ananatis* (mutant) | [18] | Yes | ATGAAGAAAACGGTAGTAATAGGCGCTGGTTTTGGAGGATTAGCTTTG<br>GCCATTCGTTTGCAGGCCGCTGGCATTCCCACCGTATTGTTAGAACAG<br>AGGGACAAGCCCGGTGGTCGTGCCTACGTCTATGAAGACCAGGGCTTC<br>ACATTCGACGCAGGACCAACGGTAATAACTGACCCTTCAGCTATAGAG<br>GAGTTATTCGCCCTTGCCGGGAAGCAATTAAAAGAATATGTTGAGCTA<br>CTGCCGGTTACACCCTTCTATAGGCTTTGTTGGGAGTCTGGCAAAGTGT<br>TCAACTATGATAACGATCAGACCCGTCTGGAAGCGCAAATCCAGCAGT<br>TTAATCCCAGGGACGTTGAAGGTTACAGACAGTTCCTTGATTACAGTA<br>GAGCTGTCTTTAAGGAAGGATACTTGAAATTAGGGACGGTCCCCTTCC<br>TATCCTTCCGTGATATGCTGCGTGCAGCTCCTCAGTTAGCTAAACTGCA<br>AGCATGGCGTTCCGTTTACTCTAAGGTAGCATCCTACATTGAAGATGA<br>GCACTTGAGACAGGCGTTCTCTTTTCACTCTCTTCTGGTAGGCGGGAAT<br>CCTTTCGCAACCAGTTCCATATACACACTTATTCACGCACTGGAACGTG<br>AGTGGGGAGTTTGGTTTCCTCGTGGGGGCACGGGTGCGTTGGTCCAAG<br>GAATGATTAAGCTGTTCCAGGATCTAGGAGGTGAAGTCGTCCTTAACG<br>CGCGTGTGTCACATATGGAAACTACTGGAAACAAAATAGAAGCGGTCC<br>ATTTGGAAGATGGCAGGAGATTCTTAACTCAGGCAGTCGCTTCCAATG<br>CGGATGTTGTTCATACTTACAGGGACTTACTTTCCCAACACCCCGCTGC<br>CGTAAAGCAATCAAACAAACTACAGACGAAGAGAATGTCTAACAGCC<br>TGTTTGTTTTATATTTCGGATTGAATCACCATCATGACCAGCTTGCACA<br>TCATACCGTATGTTTTGGTCCAAGATACAGGGAATTAATCGATGAGAT<br>ATTCAATCACGACGGACTTGCTGAAGACTTTAGTTTATACCTTCATGCT<br>CCCTGCGTGACCGATAGTAGTTTAGCGCCAGAGGGTTGTGGGTCATAT<br>TACGTGTTGGCTCCGGTACCACACCTAGGAACAGCAAATCTGGACTGG |

| | | | | |
|---|---|---|---|---|
| | | | | ACAGTGGAGGGTCCTAAACTACGTGACCGTATCTTTGCTTATTTAGAA<br>CAGCATTATATGCCCGGGTTACGTAGTCAATTGGTGACACATCGTATG<br>TTTACGCCCTTTGATTTTCGTGACCAATTGAATGCTTACCACGGCTCCG<br>CTTTTTCCGTTGAGCCGGTTCTAACACAGTCCGCATGGTTCAGGCCCCA<br>CAATCGTGATAAAACGATCACCAACCTTTATCTAGTAGGTGCAGGCAC<br>ACACCCGGGTGCGGGCATACCCGGCGTTATCGGCAGTGCAAAGGCGA<br>CCGCGGGTCTTATGCTAGAAGATCTAATCGGCTGA |
| crtY2 | *Pantoea ananatis* (mutant) | [12] | Yes | ATGCAGCCTCATTACGATTTAATTCTGGTCGGGGCAGGGCTGGCCAAT<br>GGGCTAATTGCCCTGAGGCTTCAGCAGCAACAACCTGACATGCGTATC<br>TTATTGATAGATGCGGCTCCTCAAGCTGGGGGCAACCACACATGGTCA<br>TTTCACCATGACGACCTGACTGAGTCTCAACACAGATGGATCGCACCG<br>CTGGTGGTTCACCATTGGCCCGATTATCAAGTACGTTTCCCGACGAGA<br>CGTCGTAAATTGAACTCTGGGTATTTCTGTATTACATCACAAAGATTTG<br>CCGAGGTTCTGCAACGTCAGTTCGGACCTCACCTTTGGATGGACACAG<br>CCGTCGCAGAAGTAAATGCTGAGTCTGTACGTCTAAAGAAAGGCCAGG<br>TTATAGGAGCGCGTGCGGTGATCGATGGACGTGGATATGCTGCTAACT<br>CAGCGCTTTCTGTGGGGTTCCAAGCGTTTATCGGCCAGGAATGGAGAC<br>TTAGTCACCCGCACGGGCTAAGTTCCCCCATTATTATGGACGCAACTGT<br>AGATCAACAGAACGGCTATAGATTCGTCTATTCCTTGCCACTGAGTCCT<br>ACGCGTCTTCTAATTGAGGATACTCATTATATTGACAATGCCACATTGG<br>ACCCGGAGTGTGCAAGGCAAAACATATGTGACTACGCTGCACAACAG<br>GGATGGCAGCTTCAGACCCTACTACGTGAGGAACAAGGCGCACTACCC<br>ATCACACTATCAGGTAATGCGGATGCGTTTTGGCAACAAAGGCCACTT<br>GCTTGCTCAGGACTGAGGGCAGGTCTATTTCATCCGACTACGGGGTAT<br>TCCCTGCCACTTGCTGTTGCAGTTGCTGACCGTTTAAGTGCACTAGATG<br>TATTCACTTCTGCGTCTATCCATCACGCAATCACTCACTTCGCTAGAGA<br>AAGGTGGCAACAACAAGGCTTTTTTAGAATGTTAAACAGAATGTTATT<br>CCTGGCAGGTCCGGCGGATTCCCATTGGCGTGTGATGCAAAGATTTTA<br>TGGGTTACCAGAGGATTTAATTGCACGTTTTTACGCCGGCAAACTGAC<br>GCTAACCGACAGGCTGAGAATATTATCTGGTAAACCTAGTGTGCCCGT<br>TTTAGCGGCGCTGCAAGCCATTATGACTACGCACCGTGGCTGA |
| crtE03M | *Xanthophyllo myces* | [19] | Yes | ATGGATTACGCGAACATCCTCACAGCAATTCCACTCGAGTTTACTCCTC<br>AGGATGATATCGTGCTCCTTGAACCGTATCATTACCTAGGAAAGAACC |

| | | | | |
|---|---|---|---|---|
| | *dendrorhous* (mutant) | | | CTGGAAAAGAAATTCGATCACAACTCATCGAGGCTTTCAACTATTGGT TGGATGTCAAGAAGGAGGATCTCGAGGTCATCCAGAACGTTGTTGGCA TGCTACATACCGCTAGCTTATTAATGGACGATGTGGAGGATTCATCGG TCCTCAGGCGTGGGTCGCCTGTGGCCCATCTAATTTACGGGATTCCGCA GACAATAAACACTGCAAACTACGTCTACTTTCTGGCTTATCAAGAGAT CTTCAAGCTTCGCCCAACACCGATACCCATGCCTGTAATTCCTCCTTCA TCTGCTTCGCTTCAATCATCCGTCTCCTCTGCATCCTCCTCCTCCTCGGC CTCGTCTGAAAACGGGGGCACGTCAACTCCTAATTCGCAGATTCCGTT CTCGAAAGATACGTATCTTGATAAAGTGATCACAGACGAGATGCTTTC CCTCCATAGAGGGCAGGGCCTGGAGCTATTCTGGAGAGATAGTCTGAC GTGTCCTAGCGAAGAGGAATATGTGAAAATGGTTCTTGGAAAGACGG GAGGTTTGTTCCGTATAGCGGTCAGATTGATGATGGCAAAGTCAGAAT GTGACATAGACTTTGTCCAGCTTGTCAACTTGATCTCAATATACTTCCA GATCAGGGATGACTATATGAACCTTCAGTCTTCTGAGTATGCCCATAA TAAGAATTTTGCAGAGGACCTCACAGAAGGGAAATTCAGTTTTCCCAC TATCCACTCGATTCATGCCAACCCCTCATCGAGACTCGTCATCAATACG TTGCAGAAGAAATCGACCTCTCCTGAGATCCTTCACCACTGTGTAAAC TACATGCGCACAGAAACCCACTCATTCGAATATACTCAGGAAGTCCTC AACACCTTGTCAGGTGCACTCGAGAGAGAACTAGGAAGGCTTCAAGG AGAGTTCGCAGAAGCTAACTCAAGGATGGATCTTGGAGACGTAGATTC GGAAGGAAGAACGGGGAAGAACGTCAAATTGGAAGCGATCCTGAAAA AGCTAGCCGATATCCCTCTGTGA |
| $crtI_{Nc}$ | *Neurospora crassa* | [20] | Yes | ATGGCGGAAACACAGCGTCCAAGGTCCGCCATTATCGTAGGAGCGGG AGCGGGGGGAATTGCGGTCGCAGCAAGGCTTGCGAAGGCTGGCGTGG ATGTTACGGTCTTAGAGAAAAATGACTTCACGGGGGGTAGATGCAGCC TAATCCATACAAAGGCAGGCTATAGGTTCGACCAAGGTCCGTCACTAC TTCTTCTGCCCGGGCTGTTCCGTGAAACATTCGAGGACCTAGGTACTAC ACTGGAACAAGAGGATGTTGAGCTACTTCAGTGCTTCCCTAATTACAA TATATGGTTCTCAGATGGTAAGAGATTTAGCCCCACCACTGATAACGC AACCATGAAAGTAGAAATAGAAAAGTGGGAAGGTCCAGATGGTTTA GAAGGTATCTATCCTGGCTGGCTGAGGGCCACCAGCATTACGAAACAT CCCTGAGACATGTGCTGCACAGAAACTTCAAGTCTATTCTGGAACTGG CAGACCCCAGACTGGTAGTAACGCTTTTAATGGCTTTACATCCGTTTGA |

| | | | | |
|---|---|---|---|---|
| | | | | AAGCATCTGGCACCGTGCAGGTAGGTATTTCAAGACCGACAGAATGCA AAGAGTTTTCACGTTTGCAACGATGTATATGGGAATGAGCCCGTTCGA CGCGCCAGCTACTTACTCTCTACTACAATACAGCGAGCTGGCTGAGGG TATATGGTATCCGAGAGGTGGTTTCCACAAAGTACTTGACGCCCTTGTT AAGATCGGGGAGAGAATGGGAGTTAAGTATAGGTTAAATACGGGAGT TTCACAAGTGTTGACGGATGGCGGCAAGAACGGAAAGAAACCGAAGG CGACTGGGGTGCAATTAGAAAATGGGGAGGTTCTGAATGCCGACTTGG TTGTAGTCAATGCCGACCTTGTCTACACTTATAACAACCTACTACCCAA GGAAATAGGAGGCATCAAAAAGTATGCAAATAAGCTTAATAATAGAA AAGCATCATGTTCTTCCATTTCATTTTACTGGAGCTTATCCGGAATGGC TAAGGAACTTGAGACACATAATATATTTTTAGCAGAAGAATACAAGGA GAGCTTTGATGCTATTTTTGAAAGGCAAGCGTTGCCGGATGACCCCAG CTTCTATATTCATGTACCTTCTCGTGTTGACCCGTCAGCGGCGCCTCCG GATAGAGATGCGGTCATTGCGTTGGTACCTGTAGGACATTTGTTGCAG AACGGACAGCCCGAGCTGGATTGGCCTACTTTGGTATCCAAAGCCAGG GCCGGTGTGTTAGCAACCATACAGGCCCGTACAGGACTTAGTCTATCT CCCTTAATCACAGAAGAGATAGTCAATACCCCCTATACTTGGGAGACA AAGTTTAACTTATCAAAGGGTGCGATCCTTGGTCTTGCACACGATTTCT TCAATGTTTTAGCATTTCGTCCACGTACCAAAGCGCAAGGAATGGATA ATGCGTACTTTGTGGGAGCCAGTACTCACCCTGGCACAGGGGTGCCTA TAGTGCTAGCGGGTGCAAAAATAACAGCCGAGCAGATCTTAGAGGAA ACTTTCCCTAAAAACACCAAGGTCCCTTGGACAACAAACGAAGAGCGT AACAGTGAAAGGATGCGTAAAGAGATGGATGAAAAAATCACTGAGGA GGGAATTATAATGAGGTCAAATTCTAGCAAACCCGGCAGACGTGGGTC TGATGCCTTCGAAGGTGCTATGGAAGTAGTCAACCTTTTATCCCAGAG AGCCTTCCCGCTTCTAGTGGCCCTAATGGGTGTCCTGTATTTCCTGTTG TTTGTTAGATAA |
| crtYB01M | *Xanthophyllomyces dendrorhous* (mutant) | [19] | Yes | ATGACGGCTCTCGCATATTACCAGATCCATCTGATCTATACTCTCCCAA TTCTTGGTCTTCTCGGCCTGCTCACTTCCCCGATTTTGACAAAATTTGA CATCTACAAAATATCGATCCTCGTATTTATTGCGTTTAGTGCAACCACA CCATGGGACTCATGGATCATCAGAAATGGCGCACGGACATATCCATCA GCGGAGAGTGGCCAAGGCGTGTTTGGAACGTTTCTAGATGTTCCATAT GAAGAGTACGCTTTCTTTGTCATTCAAACCGTAATCACCGGCTTGGTCT |

| | | | | |
|---|---|---|---|---|
| | | | | ACGTCTTGGCAACTAGGCACCTTCTCCCATCTCTCGCGCTTCCCAAGAC<br>TAGATCGTCCGCCCTTTCTCTCGCGCTCAAGGCGCTCATCCCTCTGCCC<br>ATTATCTACCTATTTACCGCTCACCCCAGCCCATCGCCCGACCCGCTCG<br>TGACAGATCACTACTTCTACATGCGGGCACTCTCCTTACTCATCACCCC<br>ACCTACCATGCTCTTGGCAGCATTATCAGGCGAATATGCTTTCGATTGG<br>AAAAGTGGCCGAGCAAAGTCAACTATTGCAGCAATCATGATCCCGACG<br>GTGTATCTGATTTGGGTAGATTATGTTGCTGTCGGTCAAGACTCTTGGT<br>CGATCAACGATGAGAAGATTGTAGGGTGGAGGCTTGGAGGTGTACTAC<br>CCATTGAGGAAGCTATGTTCTTCTTACTGACGAATCTAATGATTGTTCT<br>GGGTCTGTCTGCCTGCGATCATACTCAGGCCCTATACCTGCTACACGGT<br>CGAACTATTTATGGCAACAAAAAGATGCCATCTTCATTTCCCCTCATTA<br>CACCGCCTGTGCTCTCCCTGTTTTTAGCAGCCGACCATACTCTTCTCA<br>GCCAAAACGTGACTTGGAACTGGCAGTCAAGTTGTTGGAGAAAAAGA<br>GCCGGAGCTTTTTGTTGCCTCGGCTGGATTTCCTAGCGAAGTTAGGGA<br>GAGGCTGGTTGGACTATACGCATTCTGCCGGGTGACTGATGATCTTAT<br>CGACTCTCCTGAAGTATCTTCCAACCCGCATGCCACAATTGACATGGTC<br>TCCGATTTTCTTACCCTACTATTTGGGCCCCCGCTACACCCTTCGCAAC<br>CTGACAAGATCCTTTCTTCGCCTTTACTTCCTCCTTCGCACCCTTCCCGA<br>CCCACGGGAATGTATCCCCTCCCGCCTCCTCCTTCGCTCTCGCCTGCCG<br>AGCTCGTTCAATTCCTTACCGAAAGGGTTCCCGTTCAATACCATTTCGC<br>CTTCAGGTTGCTCGCTAAGTTGCAAGGGCTGATCCCTCGATACCCACTC<br>GACGAACTCCTTAGAGGATACACCACTGATCTTATCTTTCCCTTATCGA<br>CAGAGGCAGTCCAGGCTCGGAAGACGCCTATCGAGACCACAGCTGAC<br>TTGCTGGACTATGGTCTATGTGTAGCAGGCTCAGTCGCCGAGCTATTG<br>GTCTATGTCTCTTGGGCAAGTGCACCAAGTCAGGTCCCTGCCACCATA<br>GAAGAAAGAGAAGCTGTGTTAGTGGCAAGCCGAGAGATGGGAACTGC<br>CCTTCAGTTGGTGAACATTGCTAGGGACATTAAAGGGGACGCAACAGA<br>AGGGAGATTTTACCTACCACTCTCATTCTTTGGTCTTCGGGATGAATCA<br>AAGCTTGCGATCCCGACTGATTGGACGGAACCTCGGCCTCAAGATTTC<br>GACAAACTCCTCAGTCTATCTCCTTCGTCCACATTACCATCTTCAAACG<br>CCTCAGAAAGCTTCCGGTTCGAATGGAAGACGTACTCGCTTCCATTAG<br>TCGCCTACGCAGAGGATCTTGCCAAACATTCTTATAAGGGAATTGACC<br>GACTTCCTACCGAGGTTCAAGCGGGAATGCGAGCGGCTTGCGCGAGCT |

| | | | | |
|---|---|---|---|---|
| | | | | ACCTACTGATCGGCCGAGAGATCAAAGTCGTTTGGAAAGGAGACGTCG<br>GAGAGAGAAGGACAGTTGCCGGATGGAGGAGAGTACGGAAAGTCTTG<br>AGTGTGGTCATGAGCGGATGGGAAGGGCAGTAA |
| $BKT_{Hl}$* | *Haematococ cus lacustris* (mutant) | [21] | Yes | ATGCACGTCGCTTCCGCACTGATGGTGGAGCAGAAAGGAAGTGAGGC<br>AGCCGCGTCTAGTCCTGATGTGCTGAGAGCCTGGGCCACCCAATATCA<br>CATGCCAAGTGAATCTTCAGACGCCGCTAGACCGGCGTTAAAACACGC<br>GTACAAACCACCCGCCTCCGATGCTAAGGGCATTACGATGGCACTGAC<br>CATAATCGGCACATGGACAGCAGTCTTCCTACATGCTATATTTCAGATC<br>CGTCTACCGACCTCAATGGATCAATTGCACTGGCTACCAGTTAGTGAG<br>GCAACAGCCCAGCTTTTAGGTGGCAGTTCCAGTTTGTTGCACATAGCG<br>GCTGTTTTCATAGTATTAGAATTCTTATATACTGGTCTGTTCATTACGA<br>CGCACGACGCAATGCACGGAACCATAGCACTGCGTCATAGACAATTGA<br>ATGACTTATTGGGTAATATTTGTATCAGCTTATATGCTTGGTTCGACTA<br>TAGTATGCTAAGAAGAAAGCATTGGGAACATCATAATCACACCGGTGA<br>AGTTGGGAAAGACCCAGACTTCCATAAGGGAAATCCCGGGCTAGTCCC<br>CTGGTTTGCATCTTTCATGTCTAGCTATATGAGTTTGTGGCAGTTTGCA<br>AGATTGGCTTGGTGGGCTGTCGTGATGCAAATGCTTGGTGCCCCGATG<br>GCCAACTTACTAGTCTTCATGGCGGCGGCACCTATCTTGTCCGCATTTA<br>GGTTGTTTTACTTTGGAACATACTTGCCCCACAAGCCGGAACCCGGCC<br>CTGCGGCAGGATCTCAGGATATGGCATGGTTTCGTGCGAAAACCTCTG<br>AGGCGTCAGATGTTATGTCCTTTTTAACATGTTACCATTTCGATCTACA<br>CTGGGAACATCATAGATGGCCGTATGCTCCTTGGTGGCAGTTGCCCCA<br>TTGTCGTCGTCTGTCCGGCAGGGGTTTGGTTCCCGCATTAGCGTAA |
| $crtZ_{Hp}$ | *Haematococ cus lacustris* | [22] | Yes | ATGCTGTCTAAATTGCAAAGCATTTCAGTAAAAGCGAGACGTGTCGAG<br>TTAGCGAGGGATATCACGAGGCCAAAAGTGTGCTTGCATGCGCAAAG<br>ATGCAGCCTAGTACGTCTTAGGGTGGCAGCACCGCAGACGGAAGAGG<br>CAGTCGGCACGCAACAAGCGGCAGGCGCTGGGGATGAACATAGCGCG<br>GATGTCGCTTTACAACAACTTGATAGGGCTATAGCCGAAAGACGTGCT<br>CGTAGAAAACGTGAGCAACTGTCTTACCAAGCGGCAGCGATAGCAGC<br>CTCCATAGGAGTCAGCGGTATTGCTATCTTCGCTACTTACTTGCGTTTC<br>GCCATGCATATGACGGTTGGAGGTGCCGTGCCCTGGGGCGAAGTTGCT<br>GGCACGTTGTTGCTTGTTGTCGGGGGTGCTTTGGGTATGGAGATGTAC<br>GCCAGATACGCTCATAAAGCCATATGGCATGAGAGCCCTTTAGGCTGG |

| | | | | |
|---|---|---|---|---|
| | | | | CTATTGCATAAGTCTCACCACACACCTAGGACGGGGCCATTTGAAGCG AATGATTTATTTGCAATCATAAATGGTCTACCGGCAATGCTACTATGCA CCTTCGGCTTCTGGCTGCCTAACGTTCTAGGGACTGCATGCTTTGGGGC GGGTCTAGGAATCACATTGTATGGCATGGCATACATGTTTGTTCACGA CGGTTTAGTACACAGGAGATTTCCCACCGGTCCGATTGCCGGGTTACC ATATATGAAACGTTTGACGGTTGCACATCAACTACACCACAGTGGGAA GTACGGTGGAGCCCCGTGGGGAATGTTTCTTGGACCACAAGAACTGCA ACATATTCCGGGGGCGGCGGAGGAAGTTGAAAGGTTAGTGCTGGAGC TGGATTGGTCAAAGAGGTAA |
| crtOx | *Staphylococcus aureus* | [23] | Yes | ATGACTAAACACATCATCGTAATTGGCGGCGGGCTGGGTGGTATTTCA GCTGCAATCAGAATGGCCCAGAGCGGATACTCTGTCTCCCTTTACGAA CAAAACACTCACATAGGGGGAAAGGTTAATAGGCATGAAAGTGATGG CTTTGGATTTGATTTAGGGCCATCTATATTAACCATGCCCTATATCTTC GAGAAGCTTTTTGAGTATTCTAAAAAGCAAATGAGCGATTACGTAACC ATTAAACGTTTACCTCACCAGTGGCGTAGCTTTTTTCCAGATGGAACCA CAATCGATCTTTACGAAGGGATTAAAGAAACTGGACAGCATAATGCGA TACTGAGCAAACAGGACATAGAGGAGCTTCAGAACTATCTGAATTACA CCAGGAGGATTGATAGAATCACTGAGAAGGGGTATTTCAACTATGGCC TGGACACCTTATCCCAAATTATCAAATTCCATGGTCCTCTGAACGCACT GATAAACTATGACTACGTACATACGATGCAACAGGCCATTGATAAGAG AATTTCTAATCCTTACTTGAGGCAGATGCTGGGCTATTTCATAAAGTAC GTAGGCTCTTCTTCCTATGATGCTCCTGCAGTACTGTCCATGTTATTTC ATATGCAACAGGAACAAGGTCTATGGTACGTAGAGGGGGGAATACAT CACCTTGCTAACGCTCTTGAGAAGCTGGCACGTGAAGAGGGCGTTACC ATACATACGGGCGCTAGAGTAGATAATATAAAGACGTATCAAAGACG TGTGACAGGAGTGAGACTTGATACAGGAGAATTCGTGAAAGCCGACT ACATAATTTCAAACATGGAGGTGATACCAACCTATAAGTACCTGATAC ACTTAGATACACAAAGATTGAACAAGTTGGAGAGAGAGTTTGAGCCA GCCAGTTCAGGATATGTGATGCACTTGGGAGTGGCCTGTCAGTACCCC CAACTGGCGCACCATAATTTCTTCTTTACAGAAAATGCCTACTTAAATT ACCAACAAGTGTTCCATGAAAAGGTATTACCTGACGATCCTACTATTT ACTTGGTGAACACAAACAAAACAGACCACACTCAGGCACCAGTTGGCT ATGAGAATATTAAAGTGTTGCCGCATATTCCGTACATTCAAGATCAGC |

| | | | | |
|---|---|---|---|---|
| | | | | CTTTTACGACCGAGGATTACGCGAAATTTCGTGACAAAATACTAGATAAACTTGAAAAAATGGGTCTGACGGATTTGAGGAAACACATTATATACGAAGATGTCTGGACACCTGAGGACATCGAAAAGAATTACAGGTCCAACCGTGGAGCCATTTACGGGGTAGTTGCAGATAAGAAGAAGAACAAGGGTTTCAAATTTCCAAAAGAATCTCAGTACTTCGAGAATTTATACTTCGTAGGAGGCTCCGTCAACCCAGGTGGCGGCATGCCGATGGTGACTTTGTCTGGACAGCAAGTGGCTGATAAGATAAATGCCAGGGAAGCTAAGAACAGGAAATAA |

### Table S6 – Sequences of CRAPS expression cassettes

| Cassette | Control elements | Locus | Sequence (5′ – 3′)[a] |
|---|---|---|---|
| 1 | $P_{FBA1}$-$T_{FUM1}$-T8-$T_{PRM9}$ | FgF20 | ATCCAACTGGCACCGCTGGCTTGAACAACAATACCAGCCTTCCAACTTCTGTAAAT<br>AACGGCGGTACGCCAGTGCCACCAGTACCGTTACCTTTCGGTATACCTCCTTTCCCC<br>ATGTTTCCAATGCCCTTCATGCCTCCAACGGCTACTATCACAAATCCTCATCAAGCT<br>GACGCAAGCCCTAAGAAATGAATAACAATACTGACAGTACTAAATAATTGCCTAC<br>TTGGCTTCACATACGTTGCATACGTCGATATAGATAATAATGATAATGACAGCAGG<br>ATTATCGTAATACGTAATAGTTGAAAATCTCAAAAATGTGTGGGTCATTACGTAAA<br>TAATGATAGGAATGGGATTCTTCTATTTTTCCTTTTTCCATTCTAGCAGCCGTCGGG<br>AAAACGTGGCATCCTCTCTTTCGGGCTCAATTGGAGTCACGCTGCCGTGAGCATCC<br>TCTCTTTCCATATCTAACAACTGAGCACGTAACCAATGGAAAAGCATGAGCTTAGC<br>GTTGCTCCAAAAAAGTATTGGATGGTTAATACCATTTGTCTGTTCTCTTCTGACTTT<br>GACTCCTCAAAAAAAAAAAATCTACAATCAACAGATCGCTTCAATTACGCCCTCAC<br>AAAAACTTTTTTCCTTCTTCTTCGCCCACGTTAAATTTTATCCCTCATGTTGTCTAAC<br>GGATTTCTGCACTTGATTTATTATAAA**AAGACAAAGACATAATACTTCTCTATCA**<br>**ATTTCAGTTATTGTTCTTCCTTGCGTTATTCTTCTGTTCTTCTTTTTCTTTTGTC**<br>**ATATATAACCATAACCAAGTAATACATATTCAAA**ACGAGCTAAATACCTAATAA<br>TATACAAGTTTTTTATGTCTTATTATATGAAGGAAAATAAGGAAGTGGAGAGAATT<br>TGTGATTCAGCAATGGTCC**TAAGTCCGCGGATAACCATT****CGGACAGAAGACGG**<br>**GAGACACTAGCACACAACTTTACCAGGCAAGGTATTTGACGCTAGCATGTGTC**<br>**CAATTCAGTGTCATTTATGATTTTTTGTAGTAGGATATAAATATATACAGCGCT**<br>**CCAAATAGTGCGGTTGCCCCAAAAACACCACGGAACCTCATCTGTTCTCGTAC**<br>**TTTGTTGTGACAAAGTAGCTCACTGCCTTATTATCACATTTTCATTATGCAACG**<br>**CTTCGGAAAATACGATGTTGAAAATAATGGAAGGTCGGGATGAGCATATACAA**<br>**GC** |
| 2 | $P_{\mathit{PGK1}}$-$T_{SYNTH11}$-T6-$T_{\mathit{HSP26}}$ | FgF24 | TCAGGCATGAACGCATCACAGACAAAATCTTCTTGACAAACGTCACAATTGATCCC<br>TCCCCATCCGTTATCACAATGACAGGTGTCATTTTGTGCTCTTATGGGACGATCCTT<br>ATTACCGCTTTCATCCGGTGATAGACCGCCACAGAGGGGCAGAGAGCAATCATCA<br>CCTGCAAACCCTTCTATACACTCACATCTACCAGTGTACGAATTGCATTCAGAAAA |

| | | | |
|---|---|---|---|
| | | | CTGTTTGCATTCAAAAATAGGTAGCATACAATTAAAACATGGCGGGCATGTATCAT<br>TGCCCTTATCTTGTGCAGTTAGACGCGAATTTTTCGAAGAAGTACCTTCAAAGAAT<br>GGGGTCTTATCTTGTTTTGCAAGTACCACTGAGCAGGATAATAATAGAAATGATAA<br>TATACTATAGTAGAGATAACGTCGATGACTTCCCATACTGTAATTGCTTTTAGTTGT<br>GTATTTTTAGTGTGCAAGTTTCTGTAAATCGATTAATTTTTTTTTCTTTCCTCTTTTT<br>ATTAACCTTAATTTTTATTTTAGATTCCTGACTTCAACTCAAGACGCACAGATATTA<br>TAACATCTGCATAATAGGCATTTGCAAGAATTACTCGTGAGTAAGGAAAGAGTGA<br>GGAACTATCGCATACCTGCATTTAAAGATGCCGATTTGGGCGCGAATCCTTTATTTT<br>GGCTTCACCCTCATACTATTATCAGGGCCAGAAAAAGGAAGTGTTTCCCTCCTTCTT<br>GAATTGATGTTACCCTCATAAAGCACGTGGCCTCTTATCGAGAAAGAAATTACCGT<br>CGCTCGTGATTTGTTTGCAAAAAGAACAAAACTGAAAAAACCCAGACACGCTCGA<br>CTTCCTGTCTTCCTATTGATTGCAGCTTCCAATTTCGTCACACAACAAGGTCCTAGC<br>GACGGCTCACAGGTTTTGTAACAAGCAATCGAAGGTTCTGGAATGGCGGGAAAGG<br>GTTTAGTACCACATGCTATGATGCCCACTGTGATCTCCAGAGCAAAGTTCGTTCGA<br>TCGTACTGTTACTCTCTCTCTTTCAAACAGAATTGTCCGAATCGTGTGACAACAACA<br>GCCTGTTCTCACACACTCTTTTCTTCTAACCAAGGGGGTGGTTTAGTTTAGTAGAAC<br>CTCGTGAAACTTACATTTACATATATATATAAA**CTTGCATAAATTGGTCAATGCAAG**<br>**AAATACATATTTGGTCTTTTCTAATTCGTAGTTTTTCAAGTTCTTAGATGCTTT**<br>**CTTTTTCTCTTTTTTACAGATCATCAAGGAAGTAATTATCTACTTTTTACAACA**<br>**AATATAAAACA**TATATAACTGTCTAGAAATAAACACCCGTCGAGCCTGTCCGATTT<br>CAAA**ACGTGTCATACGAGGTATAGCGGAGTGACCTGGCTCTTAGTGTTGTCCC**<br>**TCTCGCGAGGACCATTGTTGCTTGCATATGGGCTTGAAACATACGGTCATCAC**<br>**ATCTGAGCGATTTTACCTCTTAGAATTAGTTTAGATATATATGAGTTGATGAAT**<br>**AAATAGTTATAAAAACTTGCTTTGGCTTCGATATATGACCGTTATTTTTGACTA**<br>**AGTTTTAACGAAGGAATCTAACCTCGTGTTGAAGTCGCCTGGTAGCCCAAAAA**<br>**TGGTGTTAACATTAACTTGCTTCAATCTTTCAAATAAGT** |
| 3 | P$_{PDC1}$-T$_{SYNTH13}$-T9-T$_{GRE3}$ | FgF18 | ACATGCGACTGGGTGAGCATATGTTCCGCTGATGTGATGTGCAAGATAAACAAGC<br>AAGACAGAAACTAACTTCTTCTTCATGTAATAAACACACCCCGCGTTTATTTACCT<br>ATCTTTAAACTTCAACACCTTATATCATAACTAATATTTCTTGAGATAAGCACACTG<br>CACCCATACCTTCCTTAAAAACGTAGCTTCCAGTTTTTGGTGGTTCTGGCTTCCTTC<br>CCGATTCCGCCCGCTAAACGCATAATTTTGTTGCCTGGTGGCATTTGCAAAATGCA<br>TAACCTATGCATTTAAAAGATTATGTATGCTCTTCTGACTTTTCGTGTGATGAGGCT<br>CGTGGAAAAAATGAATAATTTATGAATTTGAGAACAATTTTGTGTTGTTACGGTAT |

| | | | |
|---|---|---|---|
| | | | TTTACTATGGAATAATCAATCAATTGAGGATTTTATGCAAATATCGTTTGAATATTT<br>TTCCGACCCTTTGAGTACTTTTCTTCATAATTGCATAATATTGTCCGCTGCCCGTTTT<br>TCTGTTAGACGGTGTCTTGATCTACTTGCTATCGTTCAACACCACCTTATTTTCTAA<br>CTATTTTTTTTTTAGCTCATTTGAATCAGCTTATGGTGATGGCACATTTTTGCATAA<br>ACCTAGCTGTCCTCGTTGAACATAGGAAAAAAAAATATATAAA**CAAGGCTCTTTC**<br>**ACTCTCCTTGGAATCAGATTTGGGTTTGTTCCCTTTATTTTCATATTTCTTGTC**<br>**ATATTCTTTTCTCAATTATTATCTTCTACTCATAACCTCACGCAAAATAACACA**<br>**GTCAAATCAATCAAA**TATATAACTGTCTAGAAATAAAGGTGCAGGCATTTCAAA**G**<br>**TAGCCCAACAGGAGCACATCGGTCCAGCCAGTAAAATCCATACTCAACGACG**<br>**ATATGAACAAATTTCCCTCATTCCGATGCTGTATATGTGTATAAATTTTTACAT**<br>**GCTCTTCTGTTTAGACACAGAACAGCTTTAAATAAAATGTTGGATATACTTTTT**<br>**CTGCCTGTGGTGTCATCCACGCTTTTAATTCATCTCTTGTATGGTTGACAAAAT**<br>**CGTCCCCAACAAAAGTGGGCTCTCAAAATTCATCACATTTAAATGCATATAGG**<br>**AAGAGCAACAGTTGGTTTGCATCTGATGTTCCTTAAAG** |
| 4 | $P_{TDH3}$-<br>$T_{SYNTH24}$-<br>T10-$T_{TPS1}$ | FgF19 | TCGAGTTTATCATTATCAATACTGCCATTTCAAAGAATACGTAAATAATTAATAGT<br>AGTGATTTTCCTAACTTTATTTAGTCAAAAAATTAGCCTTTTAATTCTGCTGTAACC<br>CGTACATGCCCAAAATAGGGGGCGGGTTACACAGAATATATAACATCGTAGGTGT<br>CTGGGTGAACAGTTTATTCCTGGCATCCACTAAATATAATGGAGCCCGCTTTTTAA<br>GCTGGCATCCAGAAAAAAAAAGAATCCCAGCACCAAAATATTGTTTTCTTCACCAA<br>CCATCAGTTCATAGGTCCATTCTCTTAGCGCAACTACAGAGAACAGGGGCACAAAC<br>AGGCAAAAAACGGGCACAACCTCAATGGAGTGATGCAACCTGCCTGGAGTAAATG<br>ATGACACAAGGCAATTGACCCACGCATGTATCTATCTCATTTTCTTACACCTTCTAT<br>TACCTTCTGCTCTCTCTGATTTGGAAAAAGCTGAAAAAAAAGGTTGAAACCAGTTC<br>CCTGAAATTATTCCCCTACTTGACTAATAAGTATATAAA**GACGGTAGGTATTGAT**<br>**TGTAATTCTGTAAATCTATTTCTTAAACTTCTTAAATTCTACTTTTATAGTTAG**<br>**TCTTTTTTTTAGTTTTAAAACACCAAGAACTTAGTTTCGAAAAACA**TGGGTGGT<br>ATGTTATATAACTGTCTAGAAATAAAGAGTATCATCTTTCAAA**TCCCCAATCGTG**<br>**GAGTGAAGCGGTGAACCCGATGCAAATGAGACGATCGTCTATTCCTGGTCCG**<br>**GTTTTCTCTGCCCTCTCTTCTATTCACTTTTTTTATACTTTATATAAAATTATAT**<br>**AAATGACATAACTGAAACGCCACACGTCCTCTCCTATTCGTTAACGCCTGTCT**<br>**GTAGCGCTGTTACTGAAGCTGCGCAAGTAGTTTTTTCACCGTATAGGCCCTCT**<br>**TTTTCTCTCTCTTTCTTTCTCTCCCGCGCTGATCTCTTCTTCGAAACAGGACCA**<br>**ACTATCATCCGCTAATTACTGACA** |

[a] Promoter and terminator homology regions used for CRAPS chromosomal repair are colored. Target sites (T#) and short synthetic terminators are bolded and underlined, respectively.
